# Hidden stop codons orchestrate mRNA fate by ambushing ribosomal frameshifting associated with codon usage

**DOI:** 10.64898/2026.01.14.699446

**Authors:** Zhanbiao Li, Xiaoqian Gu, Shimin Gong, Zeyu Wang, Qin Bian, Ziqing Wang, Jingyang Li, Shunkai Chen, Fei Li, Dan Wang, Xiuwen Li, Chuqiao Han, Xiao Liu, Qun He, Guiping Ren, Fan Lai, Zhipeng Zhou, Yunkun Dang

## Abstract

In eukaryotes, the mRNA stability is generally shaped by codon usage bias, the uneven preferences for synonymous codons, in a translation-dependent manner. However, the conserved mechanism linking codon to mRNA decay remains elusive. Hidden stop codons (HSCs), defined as stop codons located in the +1 or -1 frame relative to canonical ORF, are widespread across all genomes but largely uncharacterized. Here, we demonstrate that in both fungi and human cells, HSCs play an conserved and mechanistic role in promoting mRNA decay by rapidly terminating out-of-frame translation promoted by nonoptimal codons, primarily through +1 ribosomal frameshifting. In the filamentous fungus *Neurospora crassa*, partially deleting HSCs via synonymous substitutions in the clock gene frequency increases mRNA stability and disrupts circadian rhythmicity. In human cells, acute depletion of translation termination factor eRF1 impairs recognition of HSCs, leading to global stabilization of mRNAs enriched in nonoptimal codons and HSCs. In both *Neurospora* and humans, these mRNAs are in part degraded through the NMD pathway via UPF1. Collectively, these findings suggest that in eukaryotes, HSCs serve as surveillance elements to monitor ribosomal frameshifting and out-of-translation promoted by nonoptimal codons, thereby initiating mRNA decay through NMD.

## Introduction

The mRNA stability plays a critical role in orchestrating gene expression. The degradation of most mRNAs begins with removal of 3’ poly(A) tails, followed by the 5’ decapping, and ultimately exonucleolytic degradation from both the 5’ and 3’ end by exonuclease and exosome, respectively. ^1^ Notably, mRNA half-lives can vary vastly from minutes to days. Although mRNA stability can be influenced by microRNA, ^2^ RNA structures, ^3^ nascent peptides, ^4^ etc., codon usage bias, also termed codon optimality, has emerged as a major determinant of the mRNA stability in a translation-dependent manner, a phenomenon conserved from bacteria to humans. ^5–13^

Codon usage bias refers to the uneven usage of synonymous codons across genes and genome, which exert effects at many levels of gene expression. ^14–18^ Optimal codons are synonymous codons that are highly preferred in the genome and their corresponding cognate tRNAs are in excess, and vice versa for nonoptimal codons. ^14,19^ Due to differences in decoding efficiency, codon usage significantly affect the speed of translation elongation. ^20–23^ Consistently, in *Saccharomyces cerevisiae*, the DEAD-box helicase Dhh1p and the N-terminal domain of Not5, a component of Ccr4-Not complex, have been shown to monitor “slow” ribosomes upon decoding nonoptimal codons, subsequently leading to mRNA degradation. ^24,25^ While DDX6, the mammalian homolog of Dhh1p, were shown to degrade inefficiently translated mRNAs in human cells, ^26^ this mechanism might not be universal. For example, in the filamentous fungus *Neurospora crassa*, *dhh1* knockout does not disrupt the link between codon usage and mRNA abundance. ^27^ Therefore, whether a conserved molecular mechanism exists in eukaryotes remains elusive.

Ribosomal frameshifting is another phenomenon implicated to link with codon usage. Ribosomal frameshifting refers to a process in which translating ribosomes switch from the canonical reading frame (0 frame) into either the +1 or -1 frame during elongation, producing *trans*-frame proteins with N-terminal sequence encoded in canonical ORF and C-terminal sequence in +1 or -1 frame. ^28^ Specifically, +1 ribosomal frameshifting (+1FS) refers to the slippage of ribosome one nucleotide toward 3’ end of the ORF, whereas -1FS is slippage one nucleotide toward 5’ end. Although spontaneous translation error occurs at very low rate (∼10^-3^ to 10^-7^/codon), certain sequence contexts can promote highly efficient frameshifting at defined positions (up to 80%), known as programmed ribosomal frameshifting (PRF). ^29^ In endogenous genes of eukaryotes, apart from the canonical +1 or -1 PRF, ^30,31^ ribosomal frameshifting was speculated to pervasively occur via tRNA slippage at the P site upon A site remain unoccupied, commonly due to scarcity of charged cognate tRNAs. ^32^

Since nonoptimal codons are associated with ribosome stalling due to limited tRNA availability, ^6,25,33,34^ early computational studies predicted a link between codon usage and ribosomal frameshifting potential. ^35^ Supporting this, rare arginine codons in human mitochondrial genome can promote ribosomal frameshifting, possibly in coordination with other cis-elements. ^36^ In cancer cells, tryptophan shortage can stimulate both +1FS and -1FS at UUG codon (encode tryptophan) to generate aberrant *trans*-frame and out-of-frame peptides. ^37,38^ Furthermore, in cancer cells tRNA modifications was found to play a role in frameshifting likely by influencing reading frame maintenance. ^39,40^ In human cells, codon repeats could also induce frameshifting, leading to *trans*-frame proteins with diverse functions. ^41–43^ In addition, during translation initiation, start codon recognition also results in pervasive ribosomal frameshifting. ^44^

Hidden stop codons (HSCs) are stop codons present in the +1 or -1 frames relative to the canonical ORF, which is commonly found in all genomes. ^45^ Given that ribosomal frameshifting may pervasively occur in eukaryotic genomes, the “ambush hypothesis” proposed that HSCs rapidly terminate these aberrant frameshifting events, thus conserving energy and prevent synthesis of potentially harmful proteins. ^45^ Computational studies indicate that HSCs locate more frequently at the downstream of “shifty” codons and under positive selection, ^46,47^ yet the biological function of HSCs remains largely unexplored.

Circadian rhythm is an internal timekeeping system that enables organisms to anticipate and adapt to the 24-hour cycle of day and night. ^48^ In the model filamentous fungus *Neurospora crassa*, the circadian oscillator centers on the FREQUENCY (FRQ) protein, which forms a complex with additional components to phosphorylate the White Collar Complex (WCC), a key transcriptional activator. ^49^ Phosphorylated WCCs dissociates from chromatins, thereby repressing *frq* and other clock-controlled gene expression. ^50^ Meanwhile, Progressive degradation of FRQ and *frq* mRNA relieves this repression, allowing *frq* transcription to resume. ^49^ This negative feedback loop underlines the core of the *Neurospora* circadian clock. The *frq* gene is characterized by poor codon usage, which impact on FRQ protein conformation and thereby the circadian rhythmicity. ^20^ In this study, we show that depletion of HSCs in *frq* impairs circadian rhythm by increasing mRNA stability, likely due to failed termination of ribosomal frameshifting induced by nonoptimal codons. At genome levels, in both fungi and human cells, we show that nonoptimal codons universally induce ribosomal frameshifting, which is terminated by adjacent HSCs, leading to mRNA degradation in part via the nonsense-mediated mRNA decay (NMD) pathway. Our findings reveal a conserved mechanism in which HSCs regulate mRNA stability by rapidly terminating ribosomal frameshifting induced by nonoptimal codons.

## Results

### Deleting hidden stop codons impairs circadian clock in *Neurospora crassa*

To explore the relationship between HSCs and codon usage in eukaryotes, for each gene (longest ORF if multiple splicing variants exist), we calculated the HSC densities (number of HSCs per kilobase within the annotated ORF) and codon bias index (CBI; in which CBI=1 indicates completely optimal codon usage and CBI<0 indicates more nonoptimal codon usage in the gene). ^51^ Codon optimality was defined according to the Codon Statistics Database. ^52^ Remarkably, in all examined organisms from yeasts to humans, we observed a consistent negative correlation between the HSC density and CBI (Figure 1A). It should be noted that these organisms have different codon usage biases. In yeasts and *Caenorhabditis elegans*, codons prefer A/U at the wobble position (GC3), whereas C/G are preferred in *N. crassa*, mice and humans. These data suggest that HSCs are associated with codon usage to play certain biological functions, thus undergo evolutionary selection independent of genome nucleotide preference.

**Figure 1.**
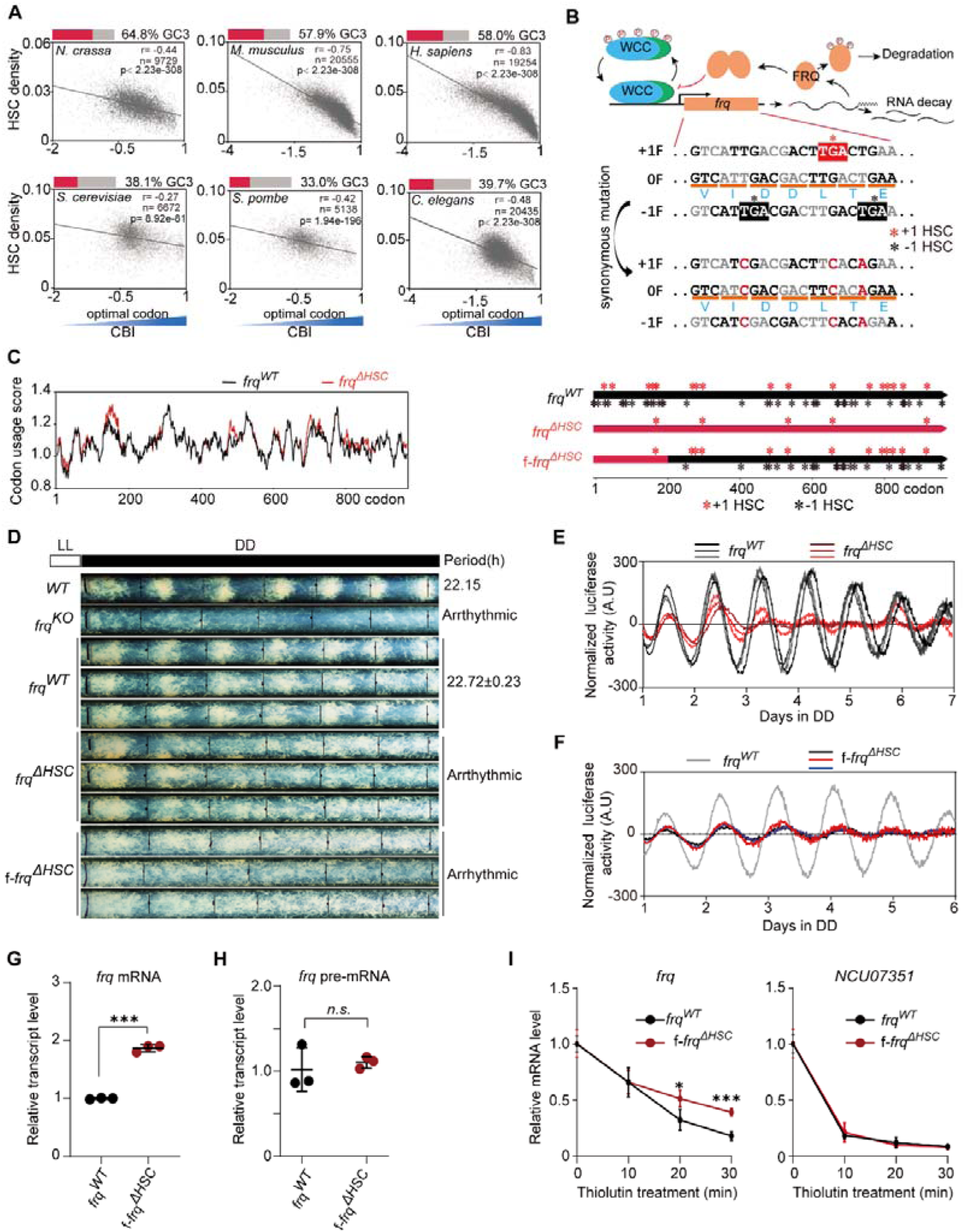
Deleting HSCs impairs circadian rhythm of *N. crassa* by affecting *frq* mRNA stability. (A) Scatter plots showing the correlation between hidden stop codon (HSC) density (number of HSCs per kb of the ORF) and the codon bias index (CBI) of genes. GC3 content of each species is depicted as the red bar. (B) Schematic diagram showing the strategy to remove HSCs with synonymous substitutions. (C) Diagrams showing the codon usage and HSC positions of wild-type (*frq^WT^*) and HSC-depleted (f-*frq*^Δ*HSC*^ and *frq*^Δ*HSC*^) sequences. Codon usage score is the mean of RSCU values in a 20-codon window. Red and black stars in the right panel indicate HSCs in +1 or -1 frame. (D) Race tube assays showing the rhythmic conidiation phenotypes in constant darkness (DD). Black lines indicating the growth fronts in every 24 h. LL, constant light. (E and F) *Luciferase* reporter driven by *frq* promoter showing the rhythm of *frq* promoter activity in *frq^WT^, frq*^Δ*HSC*^ and f-*frq*^Δ*HSC*^ strains in DD conditions, respectively. Each line indicates an individual strain. (G and H) Strand-specific RT-qPCR illustrates *frq* transcription level mRNA and pre-mRNA in *frq^WT^* and f-*frq*^Δ*HSC*^ strains under LL condition. β*-tubulin* was used as the internal control. (I) RNA decay assay for *frq^WT^* and f-*frq*^Δ*HSC*^ strains treated with thiolutin. The transcript level was normalized by spike-in human *GAPDH* gene. *NCU07351* gene was used as an internal control. For (G-I), error bars for standard deviations of three independent biological replicates. *P* values were calculated by unpaired two-tailed *t*-test (n = 3 biological replicates). *, *P* < 0.05. **, *P* < 0.01. ***, *P* < 0.001. *n.s.*, no significant.

To investigate the biological role of HSCs, we selected the *frq* gene of *Neurospora*, which centers to the circadian oscillator (Figure 1B). ^49^ The *frq* ORF harbors 61 HSCs and is known have poor codon usage. ^20^ We generated a *frq* mutant by deleting 56 HSCs via synonymous codon substitution (*frq*^Δ*HSC*^) (Figure 1B and 1C), while preserving a comparable pattern of codon usage bias as that of the wild-type *frq* (*frq^WT^*) (Figure 1C). We individually introduced the *frq^WT^* and *frq*^Δ*HSC*^ constructs into *frq* null strain (*frq^KO^*), and examined the circadian rhythm with race tube assays in the constant darkness (DD) condition. ^53^ To our surprise, conidiation rhythms in independent *frq*^Δ*HSC*^ transformants were severely impaired, becoming completely arrhythmic after 48 hours (Figure 1D). Consistently, we found that the rhythmic expression of both *frq* mRNA and FRQ protein in *frq*^Δ*HSC*^ strains were dampened compared to those in *frq^WT^* strains (Figure S1A and S1B). Furthermore, by introducing a luciferase reporter driven by the *frq* promoter (*Pfrq-luc*) in those strains as previously described, ^54^ we observed that rhythmic *frq* promoter activities was also markedly attenuated in *frq*^Δ*HSC*^ strains (Figure 1E). Together, these results indicated that HSCs in the *frq* gene may act as an important *cis*-element to maintain circadian rhythmicity in *N. crassa*.

### HSCs in 5’ region of *frq* are required for maintaining circadian rhythmicity

To evaluate the regional effect of HSCs within the *frq* ORF, we selectively deleted HSCs from the 5’ (n-*frq*^Δ*HSC*^), middle (m-*frq*^Δ*HSC*^), or 3’ (c-*frq*^Δ*HSC*^) region of the *frq* ORF (Figure S1C). Notably, only deleting the 5’ HSCs significantly impaired the circadian rhythm at both physiolocial and molecular levels (Figure S1C-E). Further dissection of the HSCs within n-*frq*^Δ*HSC*^ sequence revealed that deletion of just the first 21 HSCs (f-*frq*^Δ*HSC*^) was sufficient to recapitulate the arrhythmic phenotypes observed in both *frq*^Δ*HSC*^ and n-*frq*^Δ*HSC*^ strains, at both physiological and molecular levels (Figures 1C-D, 1F, S1B and S1F-G).

Synonymous codon optimization in *frq* coding region has been documented to affect circadian rhythm by altering the protein folding. ^20^ To exclude the possibility that this phenotype was due to a modest increase of codon usage rather than the loss of HSCs, we created a n-*frq^mOPT^*stain by optimized 15 codons to match the codon usage pattern of n-*frq*^Δ*HSC*^ sequence, while preserving all original HSCs (Figure S1H). The resulted n-*frq^mOPT^* stain exhibited normal rhythmicity, comparable to the wild type (Figure S1H). In addition, by performing the polysome profiling, we observed no significant difference of *frq* mRNA distribution in monosome and polysome fractions between *frq^WT^* and f-*frq*^Δ*HSC*^ strains, indicating that the modest increase of codon usage in f-*frq*^Δ*HSC*^ has minimal influence on translation efficiency (Figure S1I). Together, these data suggest that the impaired circadian rhythmicity observed in the f-*frq*^Δ*HSC*^ and n-*frq*^Δ*HSC*^ strains is largely due to the loss of HSC in the 5’ end region of the *frq* ORF.

### HSC depletion impairs circadian rhythm by altering *frq* mRNA stability

To explore how HSCs impact circadian clock function, we compared *frq* expression among various mutant strains. Under constant light (LL) condition, f-*frq*^Δ*HSC*^ strains showed a significant increase in FRQ protein levels compared to *frq^WT^* strains (Figure S2A). Because extensive codon-optimization of central clock gene disrupts circadian rhythmicity by altering protein structure and protein-protien interactions, ^20,55^ we ask if minor change of codon usage in f-*frq*^Δ*HSC*^ may also alter FRQ protein structure and function.

To address this, we assessed the FRQ protein stability and conformation state with three independent assays: translation inhibition, trypsin sensitivity and sensitivity to freeze-thaw cycles. ^20^ All three assays showed no significant change between f-*frq*^Δ*HSC*^ and *frq^WT^* strains (Figure S2B-D). Furthermore, by using FRQ antibodies to co-immunoprecipitation (co-IP) White Collar 2 (WC-2, the transcription activator known to interact with FRQ^56^), we observed no significant enrichment of WC-2 in f-*frq*^Δ*HSC*^ strains compared to *frq^WT^* strains (Figure S2E). These data suggesting that minor increase of codon usage due to HSC mutations has minimal influence on FRQ protein structure in f-*frq*^Δ*HSC*^ strains.

At the mRNA levels, we observed a nearly 2-fold increase of the steady-state *frq* mRNA level in the f-*frq*^Δ*HSC*^ strains compared to *frq^WT^* strains (Figure 1G). However, levels of *frq* precursor mRNA (pre-mRNA) were similar between these strains (Figure 1H), suggesting that the increased *frq* mRNA levels in f-*frq*^Δ*HSC*^ strain may be resulted from increased *frq* mRNA stability rather than enhanced transcription. Supporting this hypothesis, we found that *frq* mRNAs in the f-*frq*^Δ*HSC*^ strain decayed more slowly than in the *frq^WT^* strain upon inhibiting transcription with thiolutin (fungal RNA polymerase inhibitor), ^57,58^ while mRNAs of 4 endogenous genes maintained similar decay rates (Figures 1I and S3A). Given the important role of normal *frq* mRNA degradation in the circadian clock, ^57^ these data suggest that HSCs impact circadian rhythms in *N. crassa* at least in part though modulating *frq* mRNA stability.

### HSCs terminate aberrant out-of-frame translation in the *frq* ORF

Early computational studies have proposed that HSCs function to terminate aberrant out-of-frame translation events. ^45^ To investigate if such events occur within the upstream region of the *frq* ORF, we employed a cell-free *in vitro* translation system to monitor the out-of-frame translation as we previously described. ^22,33^ We created a set of chimeric mRNAs by fusing the f-WT or f-Δ*HSC* sequence (1-600nt of *frq^WT^* or *frq*^Δ*HSC*^) with downstream *FLuc* positioned in 0, +1 or -1 frame, respectively (Figure 2A, upper panel). As the f-Δ*HSC* lacks stop codon at any frame, the FLuc expressed from +1 or -1 frame serves as the readout for aberrant out-of-frame translation.

**Figure 2.**
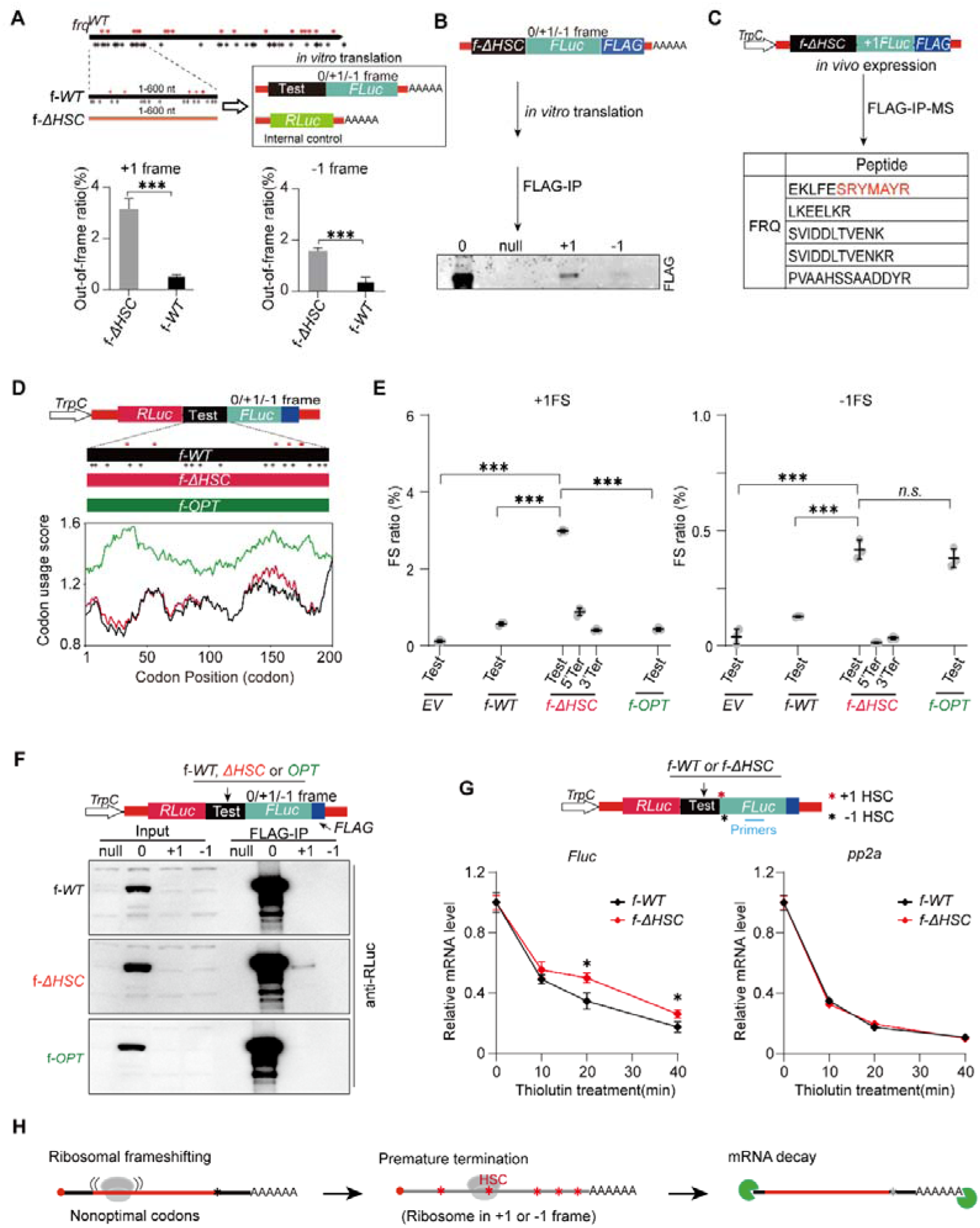
Ribosomal frameshifting is induced by nonoptimal codons in the *frq* ORF. (A) Detection of out-of-frame translation in *frq* upstream region using *in vitro* translation assay. Upper panel, the information of test sequences (f-*WT* and f-Δ*HSC*) and related reporter genes. Equal amount of *RLuc* mRNA was used as internal control. (B) Western blot analyses detecting the out-of-frame proteins derived from f-Δ*HSC* sequence by FLAG immunoprecipitation of *in vitro* protein expression. 0, +1, -1 indicates *FLuc-FLAG* is placed in corresponding frames. Null, no mRNA added. (C) Mass spectrometry identifying *in vivo* expressed *trans*-frame product. The construct is driven by a constitutive *TrpC* promoter. Black and red peptide sequences indicate amino acid residue encoded by 0 or +1 frame, respectively. The entire detected peptides were listed in Figure S4A. (D) Schematic diagram of the *in vivo* dual luciferase system and the codon usage of corresponding *frq* sequences. * indicates HSC distributions, with red and black in +1 and -1 frame. (E) Frameshifting ratios determined by dual luciferase assay as depicted in panel (D). Each line indicates the result from three independent strains. EV, empty vector. (F) Western blot representing FLAG-IP results for detecting ribosomal frameshifting in *frq* sequences (f-*WT,* f*-*Δ*HSC* and f-*OPT*) *in vivo*. (G) RNA decay assay for strains containing the dual luciferase constructs with f-*WT* or f-Δ*HSC* insertion (panel D). The transcript level was normalized with spike-in human *GAPDH* gene. *pp2a* was used as an internal control. (H) Schematic diagram of working model. For (A, E and G), error bars representing standard deviation of three independent biological replicates. *P* values were calculated by unpaired two-tailed *t-*test. ***, *P*< 0.001. **, *P*< 0.01. *, *P*< 0.05. *n.s.*, no significant.

We individually incubated these mRNAs with the cell lysates and quantified the efficiency of out-of-frame translations (out-of-frame ratio) by monitoring luciferase activities. To strictly evaluate the efficiency, luciferase signals from +1 or -1 frame were normalized with the in-frame control harboring respective test sequence. The f-Δ*HSC* sequence manifested 3.14% and 1.58% for +1 and -1 out-of-frame translation ratios, respectively (Figure 2A). Notably, these out-of-frame translation signals were much lower in the wild-type sequences (f-*WT*) (Figure 2A), consistent with a role of HSCs in terminating out-of-frame translation. To further validating these findings, we appended a *FLAG* tag to the C-terminus of *luciferase* gene and performed immunoprecipitation (IP) with anti-FLAG antibodies (Figure 2B). As expected, we observed strong FLAG signals from the control (*Fluc*-*FLAG* in the 0 frame). Importantly, robust FLAG signals were also observed with the *FLuc-FLAG* placed in +1 or -1 frame, with +1 signals stronger than -1 ones (Figure 2B). Together, these data suggest that aberrant out-of-frame translation events occur during *frq* mRNA translation and are terminated by HSCs in this *in vitro* assay.

To determine whether aberrant out-of-frame translation events occur *in vivo*, we individually introduced the same reporter sequences into wild type *N. crassa* strains under native expression conditions. The resulting transformants were subjected to FLAG IP followed by mass spectrometry analysis (IP-MS) to identify the aberrant translation products (Figure 2C). Consistent with the *in vitro* data, we identified 19 and 5 peptides derived from *FLuc* located in +1 and -1 frame (Figures 2C, and S4A-B), respectively. Importantly, we identified one *trans*-frame peptide composed of a N-terminal segment encoded in 0 frame (letters in black) and C-terminal segment encoded by +1 frame (letters in red) (Figure 2C), indicative of a ribosomal frameshifting product. Collectively, these results support that the out-of-frame translation occurs in *frq* mRNAs both *in vivo* and *in vitro*, likely through ribosomal frameshifting.

### Nonoptimal codon usage in *frq* sequence mediates +1 ribosomal frameshifting *in vivo*

To determine whether the out-of-frame translation in the f-Δ*HSC* sequence arisen from ribosomal frameshifting, we employed a improved dual luciferase reporter assay as we previously described. ^43^ In the reporter, the test sequence (f-Δ*HSC* or f-*WT*) were placed in the 0 frame directly downstream of *Renilla luciferase* (*RLuc*), while the downstream *FLuc* in 0, -1 or +1 frame, respectively (Figures 2D and S4C). To eliminate potential out-of-frame translation occurred within the *Rluc* region, we inserted 2 HSCs upstream of the test sequences, in both -1 and +1 frame. ^43,59^

Insertion of the f-Δ*HSC* sequence yielded robust +1 ribosomal frameshifting (+1FS) signals (2.99%), in stark contrast to the background level of +1FS (0.12%) from the empty vector (EV) (Figure 2E). Introducing a 0-frame stop codon immediately upstream of the f-Δ*HSC* insertion (5’Ter) abolished the +1FS signal (Figure 2E), indicating that the detected +1FS event is not due to internal translation initiations. ^59^ Furthermore, introducing a +1 frame stop codon at the 3’ end of the f-Δ*HSC* insertion (3’Ter) also abolished the +1FS signals (Figure 2E), ruling out the possible splicing artifacts. ^59^ Notably, the f-*WT* insertion showed a significantly low +1FS rate (0.57%), suggesting that HSCs effectively terminate +1FS *in vivo* (Figure 2E). Moderate level of - 1FS (0.42%) were also detected in the f-Δ*HSC* sequence, albeit lower than that from the *in vitro* assays (1.58%) (Figure 2E). The -1FS signals were nearly abolished with the 5’Ter (0.01%) and 3’Ter (0.03%) modifications, indicating that -1FS also occurs but with much lower efficiency than the +1FS (Figure 2E). Collectively, these data suggest that most out-of-frame translation events in the f-*frq* region are derived from ribosomal frameshifting, which are efficiently terminated by HSCs *in vivo*.

To assess if codon usage affects frameshifting efficiency, we designed a codon-optimized f-Δ*HSC* sequence lacking any HSC in +1 and -1 frame, termed f-*OPT* (Figure 2D, shown in green). Remarkably, codon optimization led to a dramatic reduction of +1FS signals (0.43%), while -1FS was largely unaffected (0.38%) (Figure 2E). Consistently, FLAG IP assays exhibited a clear +1FS product with expected molecular weight in the f-Δ*HSC* construct, but no visible band in either - 1FS f-Δ*HSC* construct or +1FS f-*OPT* construct (Figure 2F). To further eliminate potential sequence effects in the dual-luciferase reporter, we swapped the positions of Rluc and Fluc in a new construct and performed FLAG IP, which yielded the same results (Figure S4D). Together, these data indicate that both +1FS and -1FS occur in the f-Δ*HSC* sequence. However, the nonoptimal codons in this sequence promote +1FS, but have minimal effect on -1FS.

### Immediate termination of ribosomal frameshifting in the *frq* sequence destabilize mRNA

To understand whether HSCs in the 600nt f-*WT* sequence contribute to the decay of the dual luciferase mRNAs by terminating frameshifting, we performed mRNA decay assay by treating those strains with transcription inhibitor thiolutin (Figure 2G, top panel). ^57^ The f-Δ*HSC* dual luciferase mRNAs were more stable than f-*WT* counterpart, while the stabilities of their endogenous mRNAs remain unchanged (Figure 2G, bottom panel), supporting the specific function of HSCs in promoting degradation of mRNAs containing ribosomal frameshifting events.

It is of note that ribosomal frameshifting arised in the f-Δ*HSC* of the dual luciferase mRNAs would be ultimately terminated by the downstream HSCs located in the *Fluc* (Figure 2G, cartoon). Similarly, for the f-*frq*^Δ*HSC*^ sequence, ribosomal frameshifting arised in 5’ region of also will also encounter downstream HSCs (Figure 1C). These data suggesting that rapid termination of frameshifting events promoted by nonoptimal codons are important to trigger *frq* mRNA decay. Consistent with this notion, preserving all original HSCs while fully optimizing *frq* in the 5’ region (f-*frq^OPT+HSC^*) also stabilized *frq* mRNAs (Figure S3F) and the thereby disrupted rhythmicity (Figure S3B). Based on these observations, we thus proposed a working model that nonoptimal codons could intrinsically promote ribosomal frameshifting, which is rapidly halted by nearby HSCs and consequently trigger mRNA decay (Figure 2H).

### Putative out-of-frame translation correlates to codon usage in *N. crassa* genome

To evaluate the genome-wide relationship between codon usage and out-of-frame translation (likely through ribosomal frameshifting) in the *Neurospora* genome, we performed high-resolution ribosomal profiling (Ribo-seq) and mapped the ribosomal footprints (RFs) onto the transcriptome (NC12) as we previously described. ^22,60^ Analyses of the A-site positions of footprints revealed a strong 3-nt periodicity (Figure S5A), with 79.28%, 16.57% and 4.15% reads located in the 0, -1 and +1 frame, respectively (Figure 3A). Recent studies revealed that at least a subset of these out-of-frame footprints (A site in +1 or -1 frame) are derived from ribosomal frameshifting ^44^, prompting us to explore whether part of the out-of-frame footprints in *Neurospora* reflect bona fide out-of-frame translation events rather than experimental noises.

**Figure 3.**
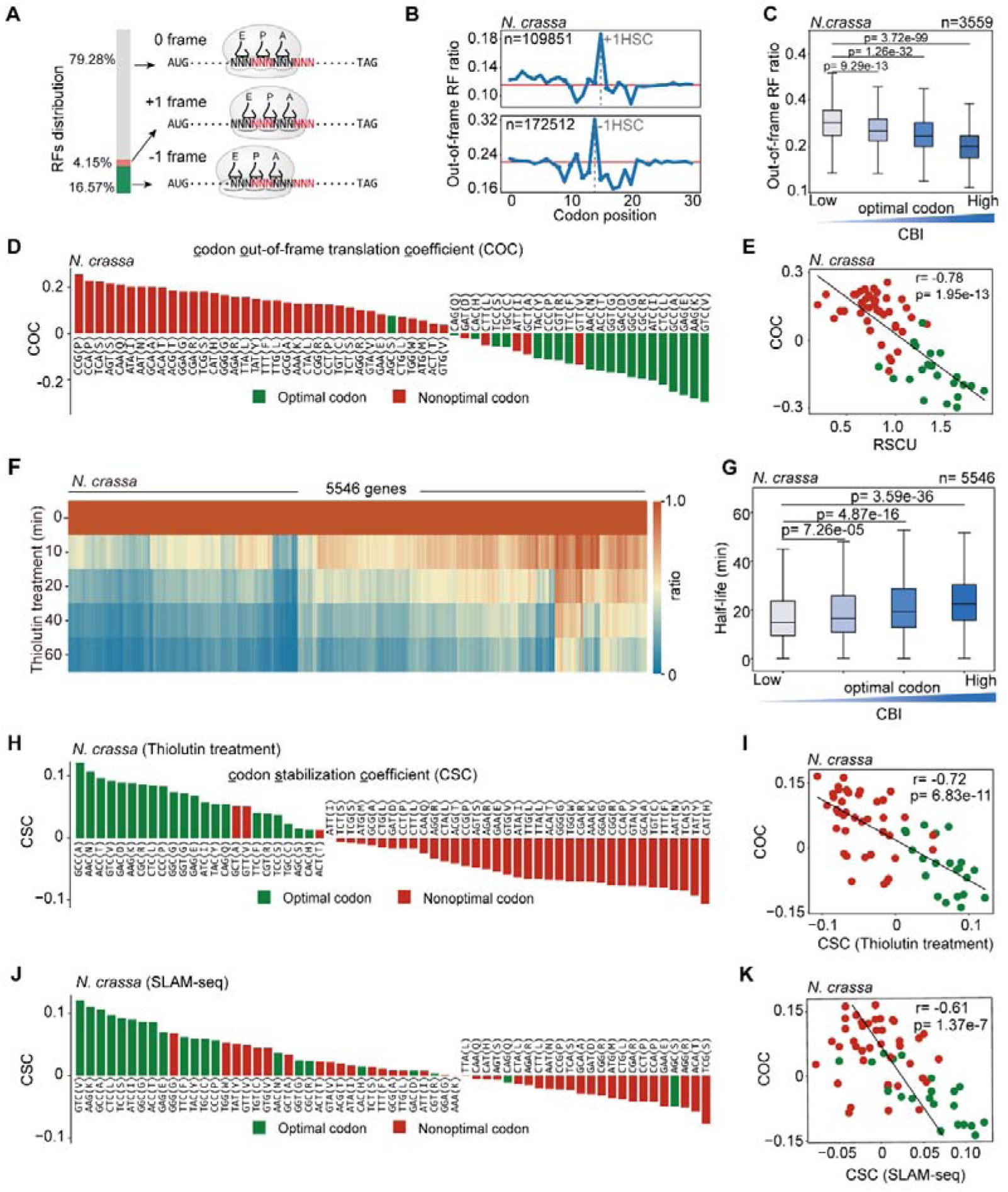
Codon usage-dependent putative out-of-frame translation is globally associated with mRNA stability in *N. crassa*. (A) schematic diagram showing the ratios of ribosome footprints (RFs) in 0, +1 and -1 frame based on the A site of each read. (B) Line plots showing the distribution of +1 or -1 out-of-frame RFs flanking corresponding HSCs. (C&G) Boxplots showing out-of-frame RF ratios (C) or mRNA half-lives (G) of genes stratified into four quantiles based on codon bias indices (CBI). n, the total number of genes. Student’s *t*-test is used for statistical analyses. (C) Bar plot displaying <u>c</u>odon <u>o</u>ut-of-frame translation <u>c</u>oefficient (COC) of each codon based on Ribo-seq data of *N. crassa*. Optimal and nonoptimal codons are marked in green and red. (D) Scatter plot showing correlation between COC and RSCU values in *N. crassa*. Optimal and nonoptimal codons are marked in green and red. (E) Heatmap showing RNA decay profiling of mRNA half-lives of 5546 genes. (H&J) Bar plots displaying codon stabilization coefficient (CSC) of each codon for RNA decay profiles based on transcription inhibition (H) or SLAM-seq (J). (I&K) Scatter plots showing correlation between COC and CSC values based on transcription inhibition (I) or SLAM-seq (K). Optimal and nonoptimal codons are marked in green and red.

Analyses of codon decoding rates (CDR) ^22^ based on the A site of each footprints revealed similar patterns across in-frame and out-of-frame codons (Figure S6), indicating the occurrence of out-of-frame translation. We next employed metagene analyses to the out-of-frame footprints, normalizing each +1/-1 RF signal with respective median RF signal of a given gene. This revealed a pronounced peak on both +1 and -1 HSCs (Figure 3B), consistent with translational pausing at stop codons. ^61^ Notably, +1 footprints exhibited an asymmetric distribution around +1 HSCs, with elevated signals upstream and diminished signals downstream, a pattern not obvious in the -1 footprints (Figure 3B). These data imply widespread occurrence of out-of-frame translation in *Neurospora* transcriptome.

To assess whether codon usage links to out-of-frame translation events, we calculated the out-of-frame footprint ratio (out-of-frame RF to all RF counts in the ORF) for each gene. As expected, we observed a robust negative correlation between out-of-frame RF ratio and CBI values, with genes enriched in nonoptimal codons have significantly higher levels of out-of-frame translation (Figure 3C). We next performed metagene analyses of +1 and -1 footprint distributions around respective HSCs in gene fragments enriched in optimal (high CBI) or nonoptimal codons (low CBI). Notably, a distinct peak of +1 footprint signals at +1 HSCs was observed in low-CBI gene fragments compared to those high-CBI ones, whereas such a difference was less pronounced at -1 HSCs (Figure S7A). These data suggest a codon optimality-dependent contribution to +1 out-of-frame translation.

Inspired by the codon stabilization coefficient (CSC) approach, ^8^ we introduced the codon out-of-frame translation coefficient (COC), which quantifies the association of each in-frame codon to out-of-frame translation potential. Specifically for each codon, we calculated the Pearson’s correlation coefficient between the out-of-frame RF ratio and the frequency of the presence of that codon in each gene (Figure 3D). Therefore, the higher the COC value, the higher chance of the codon to be associated with out-of-frame translation.

To assess the relationship between codon optimality and out-of-frame translation, we obtained the characteristics of codon optimality for each codon of *N. crassa* from the Codon Statistics Database. ^52^ Relative synonymous codon usage (RSCU) is a quantitative feature of codon optimality, which is defined as the ratio of the frequency of a given synonymous codon to expected frequency under equal usage. ^15^ Synonymous codons significantly enriched in genes with highest ENC (effective number of codons) value are defined as optimal, while the rest are considered nonoptimal. ^52^ Strikingly, most “frameshifting” codons (COC>0) coincide with nonoptimal codons, and vice versa (Figure 3D). In addition, COC and RSCU values are tightly correlated (r=-0.78) (Figure 3E), reinforcing the notion that codon optimality negatively links the frameshifting potential.

Finally, we reanalyzed an published high-resolution Ribo-seq dataset^22^ (Figure S5A), which revealed a similar tight correlation between codon usage and putative out-of-frame translation (Figure S7B-D). Notably, the COC values derived from our studies were highly concordant with those from published Ribo-seq data (r=0.98) (Figure S7E). Collectively, these data suggest that nonoptimal codons may globally mediate out-of-frame translation events in *N. crassa*.

### Putative out-of-frame translation negatively correlates to the mRNA stability in *N. crassa*

Codon usage has been well-documented as a major determinant of mRNA stability in eukaryotes. ^62^ To understand whether the out-of-frame translation may serve as a mechanistic link between codon usage and mRNA stability, we investigated the mRNA decay profile of *N. crassa* genome. We inhibited transcription with thiolutin, harvested cells at defined time points and carried out mRNA-seq. This approach enabled us to calculate the mRNA half-lives of 5546 genes (Figure 3F). We stratified these genes based on the codon bias indices (CBI) and compared the mRNA half-lives across quantiles. As expected, mRNAs enriched in nonoptimal codons manifested significantly shorter half-lives than more optimal ones (Figure 3G), consistent with previous findings that nonoptimal codon usage destabilizes mRNAs. ^8^

To assess the contribution of each codon to the mRNA stability, we computed the codon stabilization coefficient (CSC) for 61 sense codons as previously described. ^8^ Briefly, codons with negative CSC values indicate that the codon tends to destabilize mRNAs, and vice versa. As anticipated, most nonoptimal codons showed negative CSC values, while most of the optimal ones showed positively values (Figure 3H). Moreover, such a pattern could also be recapitulated with published RNA decay data (Figure S7F-G). ^63^ To rule out potential confounding effects from upstream open reading frames (uORFs), which are known to influence mRNA turnover ^64,65^, we separately analyzed genes lacking annotated uORFs as defined in Ribo-uORF^66^. These genes exhibited patterns similar to those observed in the genome-wide analyses (Figure S8A-B), suggesting that the observed correlations were not an artifact of uORF-mediated regulation.

Strikingly, we observed a strong negative correlation between CSC and COC values (r=-0.72) (Figure 3I and S8C), implying that codon usage-dependent mRNA decay might be mediated through out-of-frame translation. To independently validate that these results were not artifacts arising from transcriptional inhibition, we employed a metabolic RNA labeling approach using SLAM-seq^67^, which tracks RNA turnover by chemically labelling nascent transcripts with 4-thiouridine (s^4^U) and allowed determining half-lives of 5948 genes (Figure S7H). Consistently, SLAM-seq results demonstrated a similar association pattern between codon usage, mRNA stability and out-of-frame translation potentials (Figures 3J-K, S7I and S8C&D). Collectively, these data suggest that codon usage globally modulate transcript stability through its influence on out-of-frame translation in *N. crassa*.

### The link between mRNA stability and out-of-frame translation is conserved in mammals

We next asked whether the relationship between codon usage, putative out-of-frame translation, and mRNA stability observed in *N. crassa* is conserved across other eukaryotes. As an initial step, we analyzed published high-resolution Ribo-seq and mRNA decay data from *S. cerevisiae*, ^8,68^ a fungal species with opposite codon usage preference to *N. crassa* (Figure 1A). Consistently, we observed a strong correlation between out-of-frame translation and codon usage (Figures S7J-L), as well as a tight correlation between CSC and COC values (r=-0.79) in *S. cerevisiae* (Figure S7M).

To extend these findings to mammals, we examined published high-resolution Ribo-seq data from mouse B16 ^69^ and human K562 cell lines ^70^ (Figure S5). In both datasets, genes with higher nonoptimal codon content (low CBI values) exhibited significantly higher out-of-frame RF ratios (Figure 4A), indicating a conserved role for nonoptimal codons in promoting out-of-frame translation. At codon levels, by calculating the COC values, we observed that in both mouse and human cells, the COC values exhibited a similarly robust correlation with codon usage as defined for human and mouse ^52^ (Figures 4B-C). These data indicate that in mammals, nonoptimal codons may also promote out-of-frame translation.

**Figure 4.**
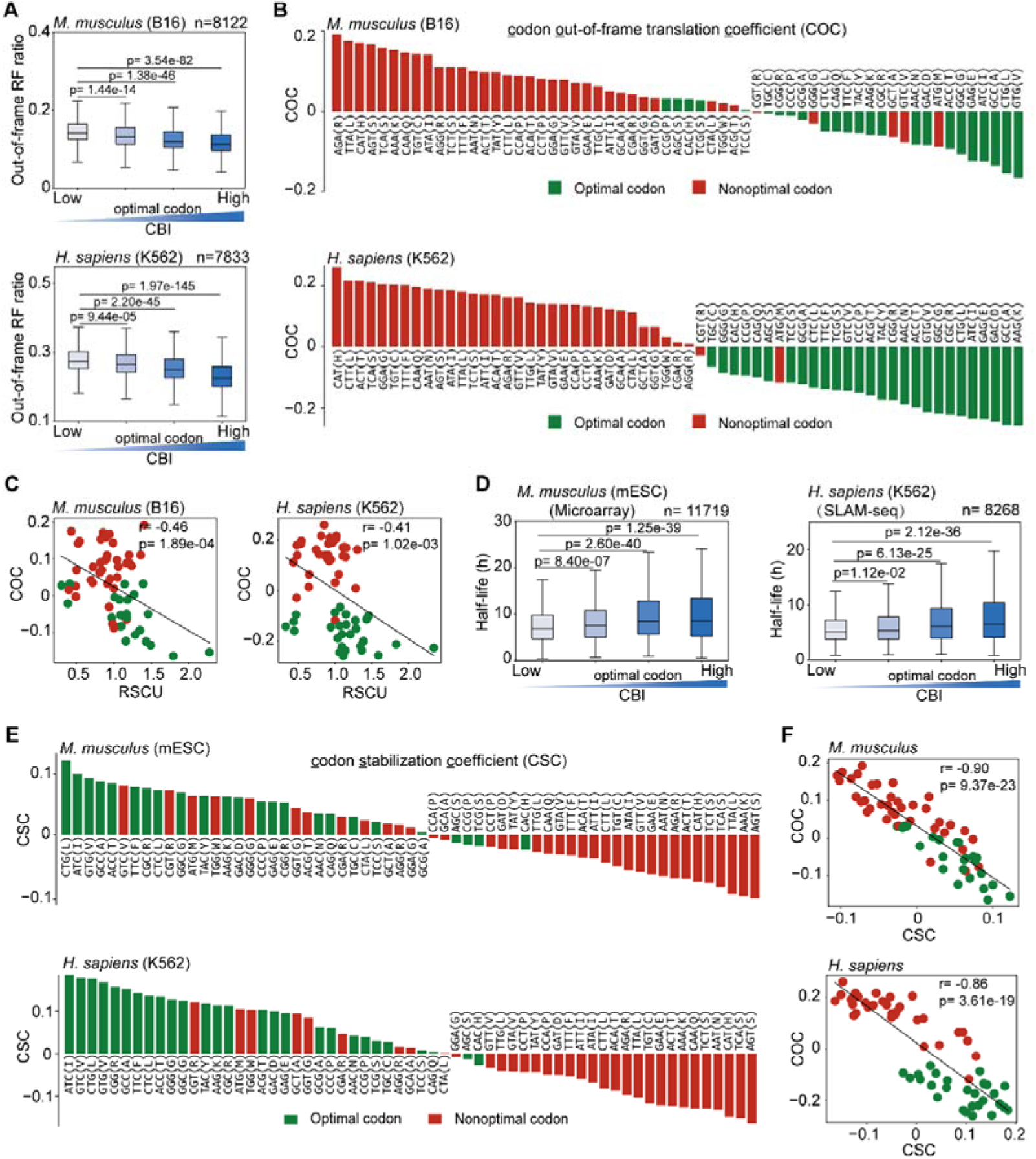
Codon usage-dependent putative out-of-frame translation globally connect with mRNA stability in mammals. (A) Boxplots showing out-of-frame ribosome footprint (RF) ratios of genes stratified into four quantiles based on CBI in *M. musculus* B16 and *H. sapiens* K562 cells. Ratios represent the mean out-of-frame RF ratio derived from two biological replicates. n, the total number of genes. Student’s *t*-test is used in A and D for statistical analyses. (B) Bar plots displaying the <u>c</u>odon <u>o</u>ut-of-frame translation <u>c</u>oefficient (COC) of each codon calculated from the mean out-of-frame translation ratios from (A) of *M. musculus* B16 and *H. sapiens* K562 cells, respectively. (C) Scatter plots showing the correlation between COC and RSCU values in *M. musculus* B16 and *H. sapiens* K562 cells. Optimal and nonoptimal codons are marked in green and red. (D) Boxplots showing mRNA stabilities of genes stratified into 4 quantiles based on CBI in *M. musculus* mESC and *H. sapiens* K562 cells. n, the total number of genes. The mRNA stability data of mESC and K562 cells were derived from the published microarray and SLAM-seq, respectively. Student’s *t*-test is used for statistical analyses. (E) Bar plots displaying <u>c</u>odon stabilization <u>c</u>oefficient (CSC) of each codon based on RNA decay data of *M. musculus* mESC and *H. sapiens* K562 cells. (F) Scatter plots showing correlation between COC and CSC values in *M. musculus* and *H. sapiens* K562 cells. Optimal and nonoptimal codons are marked in green and red.

We next assessed whether this codon-level effect on out-of-frame translation is linked to mRNA stability in mammals. We analyzed published mRNA decay profiling based on microarray of mouse embryonic stem cells (mESCs) and SLAM-seq results of human K562 cells, respectively. ^5,71^ In both cases, genes with low CBI values have significantly shorter mRNA half-lives than those with high CBI values (Figure 4D). At codon levels, we also observed the positive correlation between codon optimality and the CSC values (Figure 4E). Meanwhile, we observed a robust negative correlation between CSC and COC values in both cases (Figure 4F). Notably, from *Neurospora* to humans, these patterns persisted in mRNAs lacking annotated uORFs as defined in Ribo-uORF ^66^ (Figures S8C&E-H and S9B), ruling out uORF-mediated confounding effects.

Importantly, the Ribo-seq and mRNA decay profiling were obtained from independent research groups, minimizing the likelihood that the observed correlation arise from shared technical bias or batch effects. In sum, these findings suggest that from fungi to humans, codon usage influences mRNA stability in part by regulating out-of-frame translation, likely through ribosomal frameshifting.

### Nonoptimal codons promote ribosomal frameshifting in human cells

To examine whether the out-of-frame footprints are in part derived from bona fide out-of-frame translation in humans, we analyzed published high resolution Ribo-seq derived from liver, brain and testis tissues ^72^ (Figure S5A). Metagene analyses of +1 and -1 footprint signals revealed a pronounced peak at corresponding HSC, consistent with ribosomal pausing at stop codons (Figure 5A). Notably, we observed asymmetric distribution of +1 footprint signals, elevated upstream and diminished downstream of +1 HSCs (Figure 5A). However, such a pattern is not obvious for -1 footprints (Figure 5A). Together, these data imply widespread occurrence of out-of-frame translation in the human cells, with +1 frame translation likely being more prevalent.

**Figure 5.**
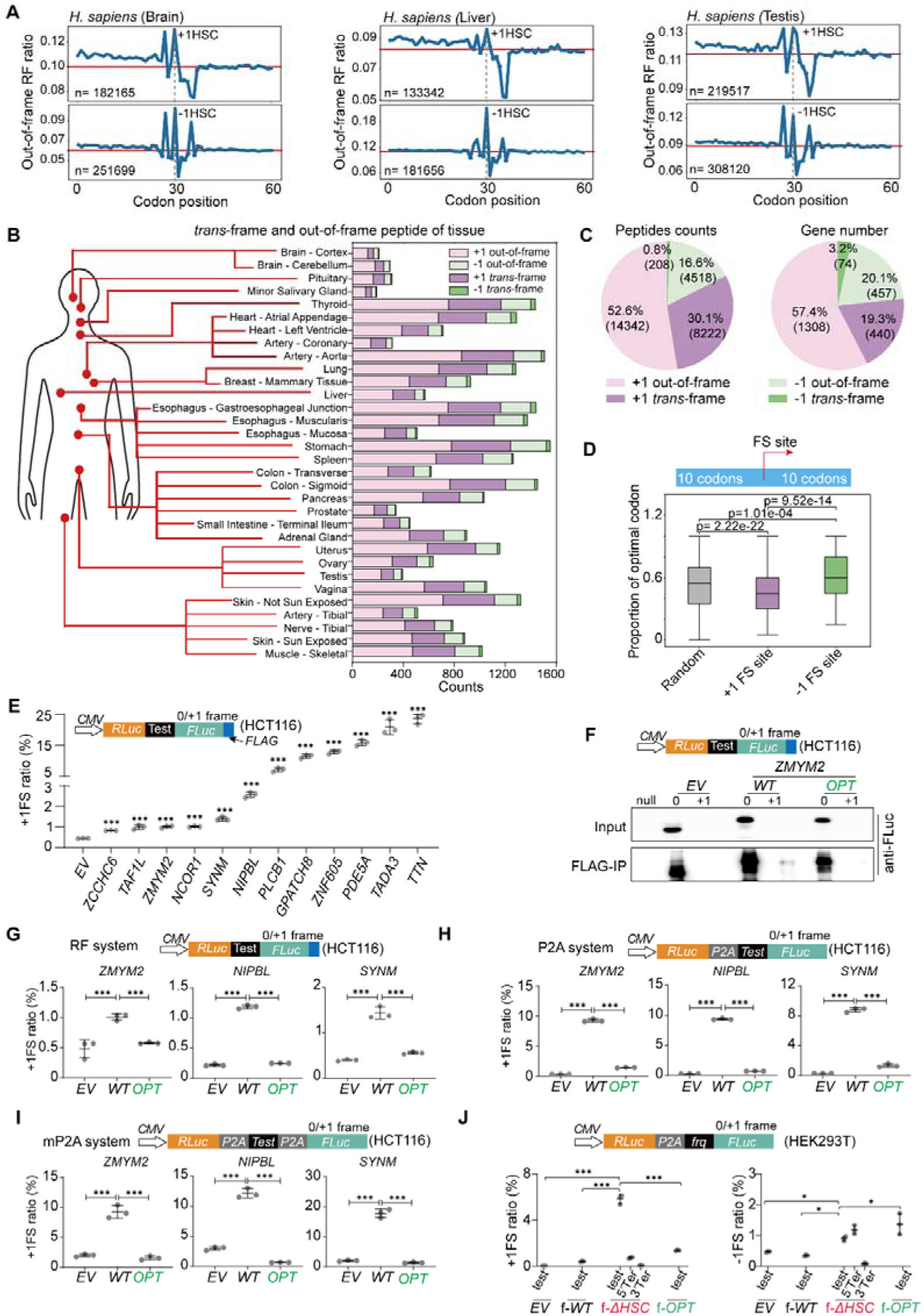
Nonoptimal codons promote ribosomal frameshifting in humans. (A) Line plots showing the distribution of +1 or -1 out-of-frame RFs flanking respective HSCs based on Ribo-seq data of *Homo sapiens* brain, liver, and testis tissue. (B) Diagram showing the *trans*-frame and out-of-frame peptides identified in published human proteome of 32 human tissues. The horizontal axis represents the numbers of identified peptides. (C) Pie charts showing the total number of identified peptides (Left) or genes harboring those peptides (Right) based on proteome data of 32 human tissues. (D) Bar plot showing optimal codon proportions within a 20-codon window of unique +1FS and - 1FS sites identified from human proteome. Random, 20-codon windows randomly selected from annotated human proteome. Student’s *t*-test is used for statistical analyses. (E) Dual-luciferase assay with HCT116 cells detecting +1 ribosomal frameshifting in 12 wild-type gene fragments (∼300 bp, sequences in Table S9) identified in human proteome. (F) Western blots showing FLAG IP of frameshift products of *ZMYM2* fragment in dual-luciferase reporter assay in HCT116 cells. *EV*, empty vector. *WT*, wild-type sequence, *OPT,* codon-optimized counterpart. (G, H and I) Detection of +1 ribosomal frameshifting in *ZMYM2*, *NIPBL* and *SYNM* fragments with three different dual luciferase reporter systems. *WT*, wild-type sequence, *OPT,* codon-optimized counterpart. EV, empty vector. The strategies of three reporter systems were shown above. (J) Dual-luciferase assay testing ribosomal frameshifting of *frq* partial sequence (Figure 2D) in HEK293T cells. Top panel, schematic diagram representing the design of dual-luciferase reporter. 5’ter, in-frame stop codon inserted upstream of test sequence. 3’ter, +1/-1 stop codon inserted downstream of test sequence. For (E and G-J), three independent repeats were performed. ***, *P*< 0.001. **, *P*< 0.01. *, *P*< 0.05. One-way ANOVA was used for significant analysis.

To examine these out-of-frame translation is in part through ribosomal frameshifting in the endogenous human genes, we analyzed a comprehensive proteome dataset containing 32 normal human tissues collected from 14 individuals^73^, using a method adapted from what we recently described for identification of ribosomal frameshifting events^43^ (Figures S10A-B). Briefly, we first scanned the entire human proteome (UniPort UP000005640) and identified 12,690 annotated proteins from both 2 biological repeats. We next generated an out-of-frame protein database based on the annotated protein sequences, integrated this database with the expressed human proteome (Table S2), then rescanned the proteome data. We further assumed that frameshifting occurs at all positions of the sequence before the identified peptides, which resulted in a database containing 61,960 putative *trans*-frame proteins (Table S3). We finally integrated this database with the annotated proteins and scanned the proteome data. To minimize false positive hits, we carefully analyzed the sequences and excluded those sequences matching unexpressed annotated proteins or microproteins (smProt). ^74^ This resulted in the detection of 27,290 peptides encoded either by alternative frame (+1/-1 out-of-frame) or by both alternative frame (+1/-1 *trans*-frame) distributed in all 32 human tissues (Figures 5B-C and S10C). These +1 or -1 out-of-frame peptides in total belongs to 1,874 and 576 unique sequences, respectively (Figure 5C (left panel) and Table S4). For +1 or -1 *trans*-frame peptides, we identified 854 and 116 unique peptide sequences, which represent 802 and 113 unique +1FS and -1FS sites within 440 and 74 genes, respectively (Figures 5C (right panel) and Tables S5-S6). The relative low levels of out-of-frame or *trans*-frame peptides might be due to the short half-lives of frameshifting products.

Analyses of genes encoding these *trans*-frame peptides revealed a striking bias in codon usage: +1FS events commonly occurred in low-CBI genes, whereas -1FS events were enriched in high-CBI genes (Figure S10D). To evaluate codon usage at the frameshifting site, we analyzed the 20-codon window flanking the 748 +1FS and 107 -1FS sites, excluding sites near termini of ORFs. As expected, +1FS sites exhibited significant low optimal codon content (proportion of nonoptimal codons to optimal ones) than the random sequence background (Figures 5D and S10E), consistent with a role for nonoptimal codons in promoting +1FS. In contrast, -1FS sites were surrounded by optimal codons (Figures 5D and S10D), suggesting a mechanistically distinct regulatory context.

To validate these findings experimentally, we randomly selected 12 sequences (∼300bp) encoding +1 *trans*-frame peptide based on the proteome data (Figure 5C), cloned the wild-type sequence fragments, transiently transfected into HCT116 cells and carried out the dual luciferase assay as in *Neurospora* (Figure 5E, table S9, sequence details). In all cases, the wild-type sequence constructs induced significant +1FS signals compared to *EV* control (Figure 5E). To confirm that the luciferase signal resulted from bona fide ribosomal frameshifting, we appended a C-terminal *FLAG* tag in the +1-frame *Fluc*, transfected into HCT116 cells and carried out IP assays. A FLAG-reactive band of expected molecular weight, comparable to the in-frame control, were detected in the *ZMYM2* +1FS construct but not in the +1 *EV* control (Figure 5F), indicating that +1FS indeed occur in the *ZMYM2* fragment.

To experimentally confirm whether nonoptimal codons promote +1FS, we codon-optimized *ZMYM2* fragments by synonymous substitutions and carried out IP assay (Figures 5F and S10F, green). The FLuc signal from *ZMYM2* +1FS construct was substantially reduced in both IP and dual luciferase assays (Figures 5F-G). Similarly, by codon-optimizing *NIPBL* and *SYNM* fragments, we also observed significant diminished +1FS signals in the dual luciferase assay (Figures 5G and S10F), implicating that nonoptimal codon usage in these sequences promotes +1FS.

Since the standard dual luciferase reporter may produce artifacts, ^59,75^ we further evaluated the 3 fragments using two independent systems proposed in recent studies, herein referred to as the P2A and mP2A system. ^43,75^ As expected, for all three wild-type sequences, these two reporter systems yielded significant +1FS signals compared to *EV* (Figure 5H-I). Consistently, in these systems, the +1FS signals were nearly abolished upon introducing the respective codon-optimized sequences (Figure 5H-I). Taken together, these data ruled out the reporter bias and confirmed the role of nonoptimal codon usage in modulating +1 frameshifting.

To further experimentally assess whether a foreign sequence harboring nonoptimal codons can also trigger frameshifting in human cells, we tested the f-Δ*HSC* sequence of *N. crassa* (Figure 2A) in human HEK293T cells with a standard dual luciferase assay. ^43,59^ It is of note that human and *N. crassa* both share strong biases for C/G-ending synonymous codons ^52^ (Table S1). Similar to that in *N. crassa*, we observed robust +1FS signals (5.84%) from the f-Δ*HSC* sequence, in contrast to the background level of +1FS from the empty vector (EV) (0.05%) or the wild-type (f-*WT*) sequence (0.43%) (Figure 5J). In addition, introducing a 0-frame stop codon before (5’Ter) or a +1 frame stop codon (3’Ter) after the f-Δ*HSC* sequence abolished the +1FS signals, supporting that the +1FS signals were not false signals derived from aberrant translation initiation or splicing ^59,75^ (Figure 5J). Moreover, the +1FS signals were dramatically decreased in the f-*OPT* construct (1.38%), indicating that nonoptimal codons are the major cause of +1FS in human cells. In contrast, the low levels of putative -1FS signals (0.91%) might be due to aberrant internal initiation and were largely independent of codon optimality (Figure 5J). Collectively, these data suggest that in human cells ribosomal frameshifting may widely occur and nonoptimal codons can promote +1FS, but might have minimal effects on -1FS in these tested conditions.

### eRF1 mediates mRNA decay by terminating out-of-frame translation in human cells

To understand whether in humans, HSCs have the similar role to regulate mRNA stability by terminating nonoptimal codon-induced frameshifting observed in *N. crassa*, we transient transfected the dual luciferase reporters harboring different version of *frq* fragments (f-*WT*, f-*HSC*, f-*OPT*) into HCT116 cells, followed by transcription inhibition with actinomycin D. Consistently, mRNAs of the f-Δ*HSC* reporter exhibited significantly higher mRNA stability than the f-*WT* reporter, while the codon-optimized reporter (f-*OPT*) showed the highest mRNA stability (Figure S3C). Collectively, these data suggest that in human cells, the HSCs contribute to mRNA decay in part by terminating nonoptimal codon-promoted ribosomal frameshifting.

In eukaryotes, eRF1 (eukaryotic release factor 1) is the only protein to recognize all stop codons, thus catalyzing the release of nascent peptides and facilitating the ribosome recycle. ^76^ Early studies revealed that premature termination of the viral-like -1 programmed ribosomal frameshifting in *CCR5* (C-C motif chemokine receptor 5) gene prompted the *CCR5* mRNA decay by stimulating the NMD pathway in mammals. ^30^ We thus hypothesized that eRF1 is responsible for recognizing HSCs upon the occurrence of ribosomal frameshifting, which subsequently trigger mRNA decay through the NMD pathway (Figure 2H).

To test this hypothesis, we employed the mini-auxin-inducible degron (mAID) system to rapidly deplete eRF1. ^77^ Specifically, the mAID tag was inserted at the C terminus of the endogenous *eRF1* loci in the HCT116:TIR1 cell line as we recently described. ^78^ After 3 hours of treatment with 10□μM 5-Ph-IAA, the targeted eRF1 proteins were nearly undetectable (Figure 6A). We next performed Ribo-seq for these cells and carried out metagene analyses, which revealed a pronounced peak at the annotated in-frame stop codons (Figure 6B), consistent with impaired translation termination. We next performed metagene analyses to +1 or -1 footprints in gene fragments stratified by codon usage (CBI). In gene fragments enriched in nonoptimal codons, the +1 footprint signals showed a asymmetric distribution, higher upstream and lower downstream of +1 HSCs (Figure 6C). Upon eRF1 depletion, the +1 footprint signals were further increased, forming a distinct peak at +1 HSCs, indicating a translation stalling at these sites (Figure 6C). In contrast, no comparable pattern was observed for -1 footprints, yet modest increase were observed near -1 HSCs (Figure S11A). These data support a model that nonoptimal codons mediates out-of-frame translation that is terminated by eRF1 at HSCs, particularly in the +1 frame.

**Figure 6.**
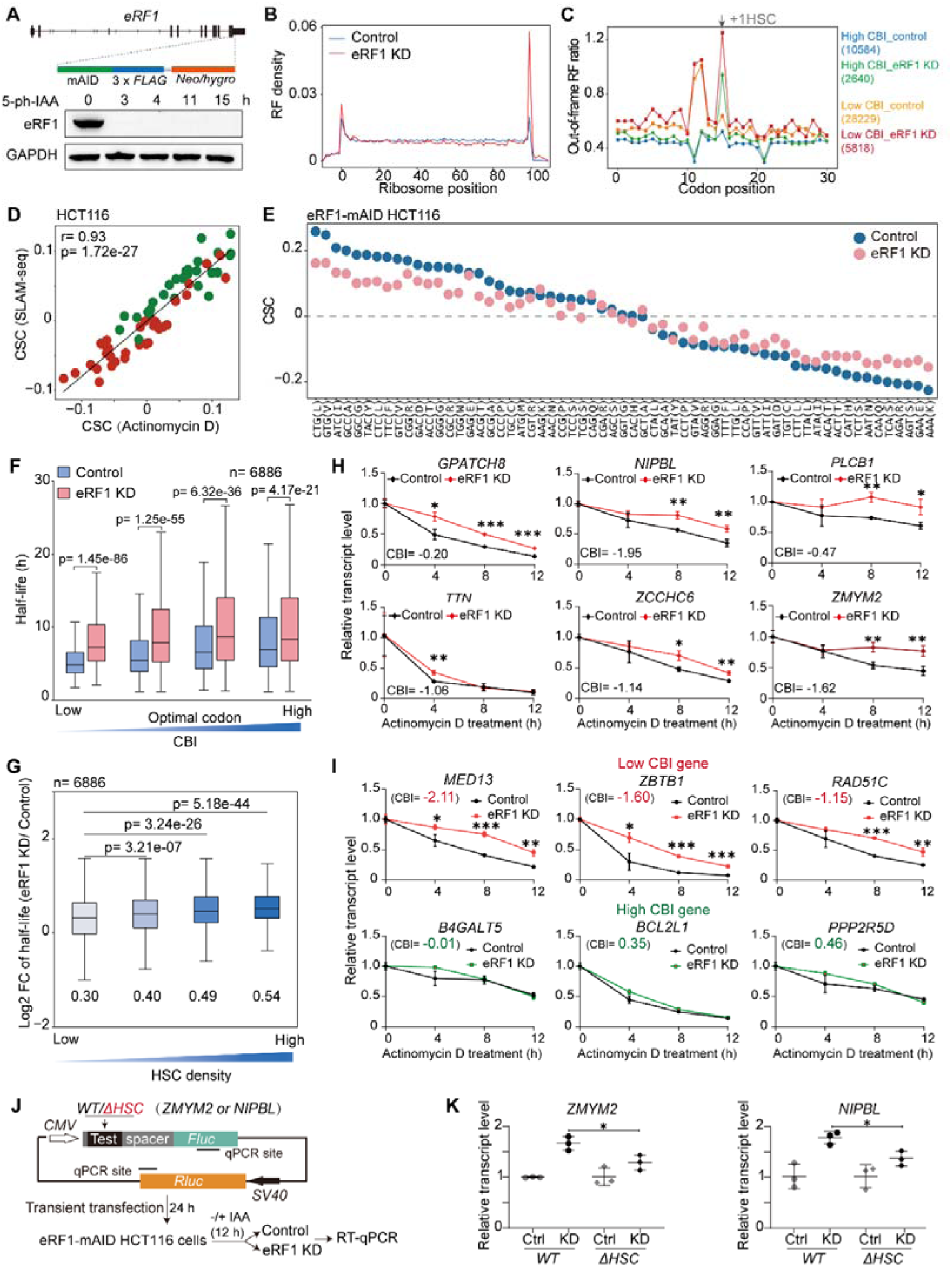
eRF1 knockdown attenuates the codon usage-mediated mRNA decay. (A) Schematic diagram (top) shows the generation of eRF1-mAID system in HCT116 cell. Western blot (bottom) shows the depletion of eRF1 after 3 hours of 5-ph-IAA treatment. (B) Metagene analysis showing ribosome footprint distribution under normal or eRF1 knock-down (KD) condition in eRF1-mAID HCT116 cells. (C) Line plot showing distribution of +1 out-of-frame ribosome footprints around +1 HSC in gene fragments with low or higher CBI, under normal or eRF1 KD condition. Fragments were stratified into 4 quantiles based on CBI, with Q1 and Q4 representing low or high CBI group, respectively. Number of fragments harboring HSC were indicated in the bracket. (D) Scatter plot demonstrating the correlation of CSC values derived from RNA decay profiles based on transcription inhibition (Actinomycin D) or SLAM-seq in eRF1-mAID HCT116 cells under normal condition. Red and green dots indicate nonoptimal and optimal codons, respectively. (E) Scatter plot of CSC values for 61 codons derived from RNA decay profiling based on transcription inhibition. The codons are arranged in descending order based on the CSC values of the control sample. Each point is marked in blue (control) or pink (eRF1 KD). (F) Boxplot displaying the effect of eRF1 KD on mRNA half-lives of genes stratified based on codon bias indices (CBI). Student’s *t*-test is used for statistical analyses. (G) Boxplot displaying the effect of eRF1 KD on mRNA half-lives of genes stratified based on HSC density. Mean of log2 fold changes of mRNA half-lives in each group is label below. Student’s *t*-test is used for statistical analyses. (H) RT-qPCR determining RNA decay profiles of 6 genes in HCT116 cells under normal or eRF1 depletion (eRF1 KD) conditions. These genes were selected from the genes harbor frameshifting site in MS result of human tissue. 3 independent repeats were performed. (I) RT-qPCR determining RNA decay profiles of 3 high CBI (top) and 3 low CBI (down) genes in RNA decay assay in HCT116 cells under normal or eRF1 KD condition. 3 independent repeats were performed. (J) Schematic diagram showing the construct and procedure to mearing abundance of mRNA containing 600nt *ZMYM2* or *NIPBL* fragments upon eRF1 KD in HCT116 cells. *WT*, wild-type sequence. Δ*HSC*, HSC-depleted counterpart. The spacer sequence lacks HSCs in +1 and -1 reading frames. (K) RT-qPCR results of expression constructs in (J). *Rluc* served as internal control. Relative mRNA levels in the control condition were set as 1. The mRNA level in eRF1 KD condition was normalized with that in control conditions. *P* values were calculated by unpaired two-tailed *t*-test (n = 3 independent samples). *, *P* < 0.05. **, *P* < 0.01. ***, *P* < 0.001.

To assess the role of eRF1 on mRNA decay, we depleted the eRF1 by add IAA for 3 hours, inhibited transcription with actinomycin D, collected samples at various time points (0, 4, 8, 12h) and performed mRNA-seq. We calculated the mRNA half-lives of each sample as described, ^79^ and determined the half-lives of 6886 genes in both control and knockdown samples (Figure S11B). Analyses of the control sample (-IAA) exhibited a similarly positive correlation between codon optimality, CSC values and half-lives (Figure S11C, left panel and S11D). Importantly, such a pattern can be recapitulated by SLAM-seq (Figure 6D and S11E).

To our surprise, depletion of eRF1 (eRF1 KD) resulted in global increases of mRNA half-lives (Figure S11B), indicating that eRF1 promotes mRNA decay. Furthermore, codon stabilization coefficient (CSC) analyses revealed a general decrease in positive CSC values (optimal codons) and increase of negative CSC values (nonoptimal codons) upon eRF1 depletion (Figures 6E and S11C, right panel), suggesting that the codon usage effect on the mRNA stability depends on eRF1. Importantly, analyses to genes lacking uORF also exhibited similar pattern (Figure S8I and S9C), minimizing the confounding effects from uORF-mediated decay.

To further confirm this effect at gene levels, we stratified the genes into four quantiles based on the CBI and analyzed their half-lives. Upon depletion of eRF1, low-CBI genes showed the most significant increase of their mRNA half-lives, while high-CBI ones showed minimal changes (Figure 6F). Moreover, genes with higher HSC densities shower higher increases in half-lives (Figure 6G). Together, these data suggest that eRF1 prefers to destabilize mRNAs harboring more nonoptimal codons and HSCs.

To validate these findings, we randomly selected 6 genes (*GPATCH8*, *NIPBL*, *PLCB1*, *TTN, ZCCHC6*, *ZMYM2*) whose fragments can trigger +1FS (Figure 5E) and carried out RT-qPCR-based mRNA decay assays with Actinomycin D. Upon eRF1 depletion, their mRNA half-lives were significantly increased (Figure 6H). We next extended this analysis to other untested genes by randomly selecting 3 genes (*MED13, ZBTB1* and *RAD51C*) with low CBI values and 3 genes (*B4GALT5, BCL2L1* and *PPP2R5D*) with high CBI values (Figure 6I), As expected, for genes with low CBI value, the mRNAs became significantly more stable upon eRF1 depletion, whereas genes with high CBI values showed very modest change of mRNA decay rate (Figure 6I).

To further confirm the role of HSCs in regulating mRNA abundance, we created reporter constructs by appending a in-frame *Fluc* to wild-type (WT) or HSC-depleted (Δ*HSC*) fragments of *ZMYM2* and *NIPBL* that can trigger +1FS (Figure 5E-I), with *Rluc* transcript as an internal control (Figure 6J). We transiently transfected these construct into HCH116 cells with or without adding IAA, inhibited with Actinomycin D for 12 hours and measured the mRNA abundance with RT-qPCR. In both cases, 12 hours after eRF1 depletion, the *WT* constructs showed a significantly higher fold-change increase in mRNA abundance than their respective Δ*HSC* counterparts (Figure 6K). It is of note that the *Fluc* sequence contains HSCs in both +1 and -1 frames (Figure 6J), which terminate frameshifting accumulated in the *ZMYM2* and *NIPBL* HSC-depleted sequences, further reinforcing the importance of immediate termination of frameshifting as observed in the *frq* sequence (Figures 2G and S3C).

Collectively, these data suggest that eRF1 plays a key role to recognize HSCs during aberrant translation, thereby linking codon usage-mediated ribosomal frameshifting and mRNA decay in human cells.

### UPF1 participates in codon usage-mediated mRNA decay in human cells

Upstream frameshift 1 (UPF1), an ATP-dependent RNA helicase, is the core factor of NMD pathway for mRNA quality control^80^. Recent studies showed that UPF1 prefer to degrade mRNAs with poorly translated coding sequences in eukaryotes^64^. Given that HSCs may act as premature termination codons (PTCs) upon ribosomal frameshifting occurs (Figure S3D), we hypothesized that UPF1 may connect with nonoptimal codons and HSCs to mediate mRNA decay in eukaryotes. At the first step, we introduced the dual luciferase construct harboring wild-type (f-*WT*) or HSC-depleted (f-Δ*HSC*) *frq* sequence fragments into *upf1^KO^* strain of *N. crassa* and performed RNA decay assay with thiolutin. Compared with that in wild-type strain (Figure 2G), no significant difference in mRNA stability were observed between f-*WT* and f-Δ*HSC* construct (Figure S12A), indicating that HSC-mediated mRNA decay is dependent on the UPF1 in this case. To confirm this effect at genome levels, we performed SLAM-seq in the *upf1^KO^* strain, which determined the mRNA half-lives of 5676 genes (Figures 7A and S12B). CSC analyses revealed that the codon optimality-dependent pattern is largely impaired in the *upf1^KO^* strain (Figure 7B and S12C), indicating a role of Upf1 in this regulatory mechanism.

**Figure 7.**
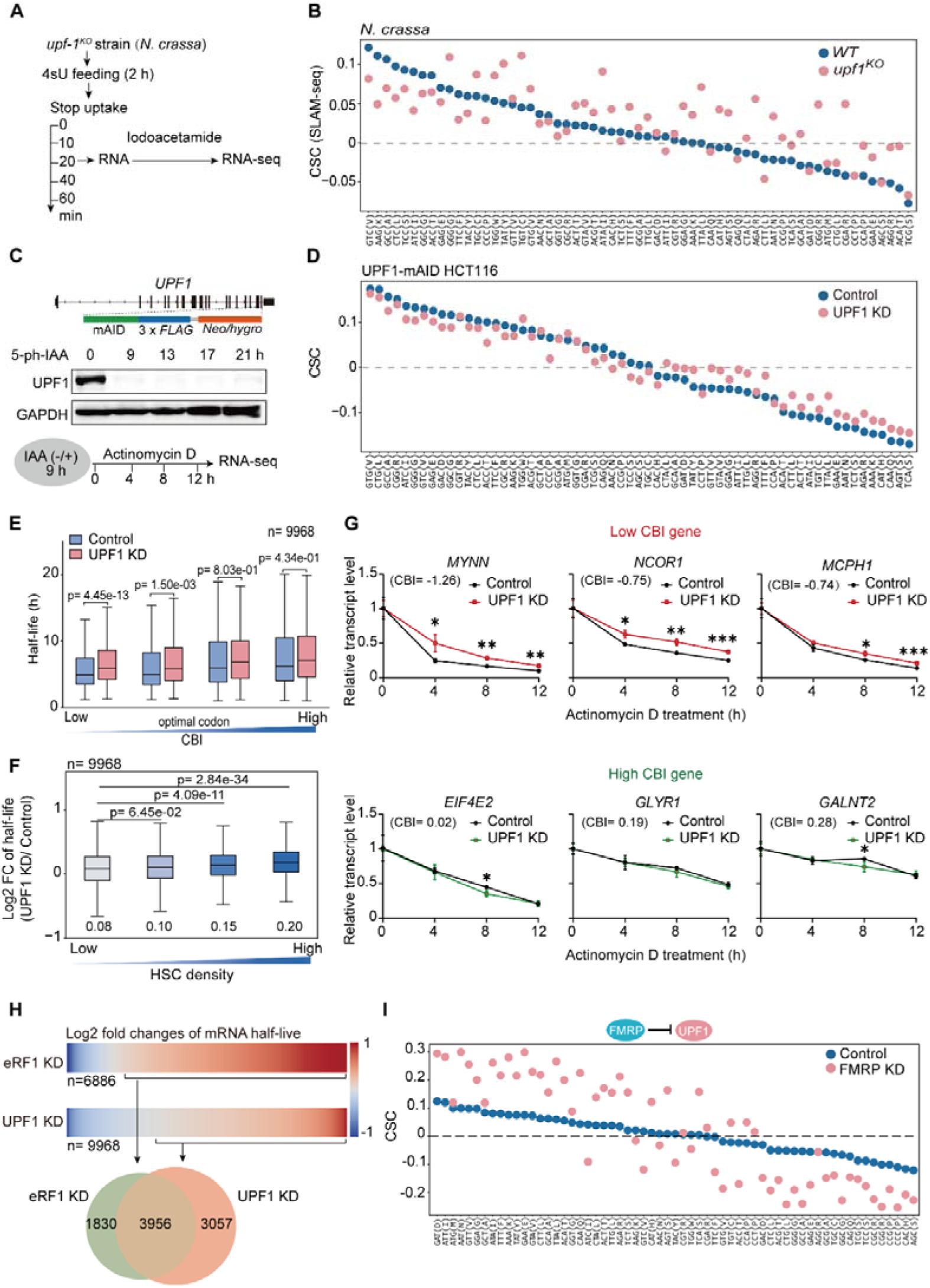
UPF1 participates in codon usage-mediated mRNA decay. (A) Schematic diagram showing the procedure of SLAM-seq assay in *upf1^KO^* strain of *N. crassa*. (B&D) Scatter plot of CSC values for 61 codons derived from SLAM-seq assay of *WT* and *upf1^KO^* strain of *N. crassa* (B) or RNA decay profile based on transcription inhibition of UPF1-mAID HCT116 cells with or without IAA (D). The codons are arranged in descending order based on the CSC values of *WT* sample. (C) Schematic diagram depicting the generation of UPF1-mAID cell line in HCT116 cell (top panel), western blot of UPF1 depletion after 5-ph-IAA treatment (middle). The design of RNA decay assay for UPF1-mAID cell shown in bottom panel. (E) Boxplot displaying the effect of UPF1 KD on mRNA half-lives across different quartiles based on gene codon bias index (CBI). Student’s *t*-test is used for statistical analyses. (F) Boxplot displaying the effect of UPF1 KD on mRNA half-lives across different quartiles based on gene HSCs density. Mean of log2 fold changes of half-lives in each group is labeled below. Student’s *t*-test is used for statistical analyses. (G) RT-qPCR results of 3 high CBI (top) and 3 low CBI (bottom) genes in RNA decay assay in HCT116 cell line upon UPF1 depletion. 3 independent replicates were performed. *P* values were calculated by unpaired two-tailed *t*-test (n = 3 independent samples). *, *P* < 0.05. **, *P* < 0.01. ***, *P* < 0.001. (H) Heatmap and Venn diagram showing the overlapped genes with increase half-lives (pie chart) between eRF1 KD and UPF1 KD cell samples. (I) Scatter plot showing the changes of CSC value of each codon upon FMRP knockdown. The analysis was based on published RNA decay data under *FMRP* siRNA in SH-SY5Y cell.

To extend this analysis to human cells, we generated an HCT116 UPF1-mAID cell line using the same auxin-inducible degron strategy as for eRF1, thereby minimizing potential secondary effects associated with stable gene knockout. Following 9 hours of treatment with 10□μM 5-Ph-IAA, UPF1 protein levels were nearly undetectable (Figure 7C). We next used Actinomycin D to inhibit transcription and quantified mRNA decay kinetics, determining half-lives for 9,986 genes in both control and UPF1-depleted samples (Figure S12D). As expected, UPF1 depletion resulted in a global stabilization of mRNAs (Figure S12D), while have minimal effects on mRNA splicing (Figure S12E). Consistent with the eRF1 depletion results, the overall codon usage effect on mRNA stability, reflected by CSC values, was markedly dampened (Figure 7D and S12F).

To further dissect the relationship between codon optimality and UPF1-dependent decay, we stratified genes into four quartiles based on their codon bias index (CBI) and analyzed changes in mRNA half-lives. As expected, genes with low CBI values—indicative of nonoptimal codon usage—exhibited a significantly greater increase in half-lives upon UPF1 depletion (Figure 7E). Consistently, genes showing the largest half-life increases were enriched for higher densities of hidden stop codons (HSCs) (Figure 7F).

Importantly, in both *Neurospora* and humans, when we specifically analyzed genes lacking upstream open reading frames (uORFs), the codon usage–dependent effects remained dampened under UPF1 depletion (Figures S8J-K), indicating that uORF translation does not confound these observations. And analyses to genes lacking uORF also exhibited similar pattern (Figure S9C), minimizing the confounding effects from uORF-mediated decay. Together, these findings suggest that UPF1 preferentially targets mRNAs harboring higher densities of nonoptimal codons and HSCs, reinforcing the role of UPF1 as a key effector in codon usage- and frameshifting-associated mRNA decay.

To confirm the deep sequencing results, we randomly selected 3 (*MYNN*, *NCOR1* and *MCPH1*) gene with low CBI values and 3 (*EIF4E2, GLYR1* and *GALNT2*) genes with high CBI values, carried out mRNA decay assays and determined mRNA levels at each time point with RT-qPCR. As expected, RT-qPCR results showed that genes harboring high levels of nonoptimal codons (CBI<0) became more stable than those with low levels of nonoptimal codons (CBI>0) (Figure 7G), further supporting the role of UPF1 in degrading mRNAs enriched in nonoptimal codons and HSCs.

To understand whether eRF1 cooperates with UPF1 to destabilize the mRNA enriched in nonoptimal codons and HSCs, we analyzed the 5786 and 7013 genes with increased half-lives from eRF1 KD and UPF1 KD cells, respectively. As expected, 68% genes with increases of half-lives upon eRF1 KD cells are coincided with those upon UPF1 KD cells (Figure 7H). Together, these data indicate that UPF1 may in part connect with eRF1 to mediate the codon usage-dependent mRNA decay.

Given the dampened codon usage effect upon depletion of UPF1, we hypothesized that the hyperactivation of the NMD pathway should, to the opposite, strengthen the effect. Fragile X mental retardation protein (FMRP), the product of *FMR1* gene, is essential for the proper synaptic plasticity and architecture, and dysfunction of FMRP causes the neurodevelopmental disorder fragile X syndrome^81^. Previous studies have shown that FMRP could directly interact with UPF1, and deletion of FMRP leads to enhanced activation of the NMD pathway^82^. Using published mRNA decay profile from FMRP KD cell lines (SH-SY5Y) ^82^, we found that the absence of FMRP strengthened the codon usage effect on mRNA stability (Figure7I). Altogether, these data suggest that UPF1, together with eRF1, contribute to codon usage-dependent mRNA decay.

## Discussion

In this study, we uncover a conserved translation surveillance mechanism, termed frameshifting stop-mediated mRNA decay (FSD), highlighting the mechanistic role of hidden stop codons in selective mRNA degradation by premature termination of ribosomal frameshifting events associated with codon usage. Our findings suggest that ribosomes translating mRNAs enriched in nonoptimal codons are more likely to shift reading frames, more frequently into the +1 than −1 frame (Figures 2 and 5). Because HSCs are more frequently embedded within nonoptimal codon-rich regions, these events increase the likelihood of premature termination and recognition by the translation termination factor eRF1, subsequently initiating mRNA degradation via the nonsense-mediated decay (NMD) pathway.

Supporting this model, genetic depletion of HSCs in the *frq* gene of *Neurospora* resulted in elevated mRNA stability and impaired circadian rhythm (Figure 1). Similarly, acute depletion of eRF1 in human cells led to a significant stabilization of mRNAs enriched in nonoptimal codons (Figure 6). Notably, even modest change in mRNA stability, such as those observed in *frq*, had profound physiological effects, emphasizing the critical role of HSCs in fine-tuning gene expression through codon usage–linked decay mechanisms (Figure 1D–F). These findings suggest that the co-evolution of HSCs and codon usage patterns has been shaped by selective pressure to maintain mRNA homeostasis in eukaryotes (Figure 1A).

Ribosomal frameshifting is universal in RNA viruses, but it was long thought to be rare events in eukaryotes. ^34^ While high-resolution Ribo-seq data revealed that out-of-frame translation events are genome-widely associated with nonoptimal codon clusters (Figures 3 and 4) ^44^, distinguishing true frameshifting events from noise has remained a challenge. Till now the dual luciferase assay remains a canonical tool for measuring ribosomal frameshifting, yet its sensitivity and specificity require rigorous controls. ^43,59,75^ In both *N. crassa* and humans, combined with multiple dual luciferase assays, immunoprecipitation and mass spectrometry, we show that the nonoptimal codons induces robust +1FS (Figures 2 and 5). Interestingly, the dual luciferase assay showed modest -1FS signals in the *frq* sequence independent of codon usage *in vivo* (Figures 2E and 5A). In line with this, genome-wide proteome data also revealed much less -1FS products, with little association with nonoptimal codon usage (Figure 5C and 5E). Interestingly, upon preparation of this manuscript, a study in budding yeasts showed that nonoptimal codon usage triggers -1FS under nutrient-limiting conditions, thus modulating mRNA abundance in part via NMD. ^83^ These findings raise the possibility that codon usage may regulate -1FS in a context-dependent manner, warranting further investigation.

HSCs have often been overlooked in mRNA surveillance. Although the “ambush hypothesis” postulated that HSCs act as checkpoints to terminate erroneous frameshifted translation, ^45,47^ their biological relevance had not been experimentally validated. Our study reveals that HSCs are evolutionarily enriched near nonoptimal codons (Figure 1A), and removal of HSCs from these regions diminishes codon usage–mediated decay, providing strong evidence for co-selection of HSCs with codon usage patterns. Notably, HSCs also exist in coding regions enriched in optimal codons, though at lower abundance. Whether such HSCs play a role in regulating mRNA stability under specific physiological conditions remains an open question. Indeed, codon optimality itself is a multifaceted concept that depends not only on genomic frequency, but also on dynamic factors such as tRNA availability, amino acid supply, and tRNA modifications. ^18,62^ Therefore, whether HSCs associated with optimal codons are also under selective pressure to mediate decay deserves further exploration.

NMD pathway was initially recognized as an mRNA quality control process by degrading faulty mRNA containing premature stop codons. ^84^ Subsequent numerous studies revealed that NMD targets to ∼10% unmutated mRNAs to regulate a variety of biological processes. ^85^ We show that depletion of eRF1 or UPF1 leads to increased mRNA stability for genes enriched in nonoptimal codons, suggesting that codon usage in part coupled with NMD pathway to dictate the fate of mRNAs via termination of ribosomal frameshifting. In line with this, a recent study showed that UPF1 has broad effect to regulate mRNA stability by targeting those mRNAs with poor translated ORFs, including ORFs with nonoptimal codon usage, in a EJC-independent manner. ^64^ Together with these findings, these data suggest that NMD is in part connects with FSD pathway to determine mRNA stability via an HSC-eRF1-UPF1 axis.

### Limitation of the study

In this study, we demonstrate a highly conserved pathway to regulate mRNA stability by terminating nonoptimal codon-induced ribosomal frameshifting through an HSC-eRF1-UPF1 axis. Although we combined dual luciferase assay with immunoprecipitation and mass spectrometry to show that nonoptimal codons could promote +1FS, a thorough investigation of ribosomal frameshifting associated with nonoptimal codons is lacking at the genome levels. Indeed, while high-resolution Ribo-seq data manifest a tight correlation between codon usage and out-of-frame translation levels at the genome level, this method is unable to distinguish ribosomal frameshifting from other aberrant translation events such as internal translation initiation. Therefore, each case has to be individually verified with canonical dual luciferase assays. Proteomic analyses provide another perspective to investigate ribosomal frameshifting at the genome level. But the sensitivity is generally much lower than the high-throughput sequencing. In addition, while we are able to identified much less but substantial amount of trans-frame peptides resulted from -1FS, these -1FS events are collectively independent of nonoptimal codon usage and the cause remains unclear.

## RESOURCE AVAILABILITY

### Lead contact

Further information and requests for resources and reagents should be directed to and will be fulfilled by the lead contact, Yunkun Dang.

### Materials availability

This study did not generate new unique reagents.

### Data and code availability

- All sequencing data that support the findings of this study have been deposited to NCBI GEO under accession numbers GSE245041, GSE238077, GSE303542 and GSE276918. The mass spectrometry data have been deposited to the ProteomeXchange Consortium via PRIDE partner repository with the dataset identifier PXD044949. Source data are provided with this paper. Published Ribo-seq data were sourced from the following accession numbers: GSE71032, GSE89704 GSE111891, GSE125218, GSE129061, PRJEB28810. Published RNA decay profiling data is available from ArrayExpress with accession number E-MTAB-7247, and NCBI with accession number SRP495209. Published proteome data is available via the ProteomeXchange identifier PXD016999.
- Custom codes used for data analysis in this paper can be found at https://github.com/GongShimin.

## ACKNOWLEDGMENTS

This work was supported by the National Natural Science Foundation of China (32470609 and 32171262 to Y.D., 32270098 and 32470073 to Z.Z., 32070626 to F. L., and 32170092 to X.L.), the Open Research Program of State Key Laboratory for Conservation (2023KF009 to Y.D.), the National Key Laboratory of Agricultural Microbiology (AML2024D02 to Z.Z), the Fundamental Research Funds for the Central Universities (2662024JC015 to Z.Z.), grants from the Yunnan Province Science and Technology Department (202401BF070001-013 to Y. D., 202201BF070001-015 to F. L., 202501AS070057 and 202401AT070436 to G.R.), grants from the Yunnan Province Education Department (2024J0004 to G.R.), and grants from The Scientific Research Fund Project of Yunnan Education Department (KC-24249887 to X. G.). We acknowledge the Dang and Lai lab members for helping many sequencing works.

## AUTHOR CONTRIBUTIONS

Y. Dang, Z. Zhou, F. Lai, Q. He, X. Liu, and G. Ren conceived and supervised the overall project. Z. Li, X. Gu and Q. Bian preformed most of the experiments. S. Gong, Z. Wang and F. Li performed bioinformatics analysis. J. Li provided helps on bioinformatics analysis. D. Wang and X. Li participated in dual-luciferase assay in human cells. Z. Wang finished rhythmic assay of *frq* protein and mRNA and co-IP assay of WC-2. C. Han and X. Liu helped to confirmed the phenotype of *frq*^Δ*HSC*^ strain on race tube. S. Chen designed *frq*^Δ*HSC*^ sequence. Q. He provided stains, antibodies, protocols and useful suggestions. Y. Dang, F. Lai, Z. Zhou, G. Ren, Z. Li, S. Gong, and X. Gu wrote the manuscript with critical feedback from all co-authors.

## SUPPLEMENTAL FIGURE TITLES AND LEGENDS

**Supplementary Figure 1.**
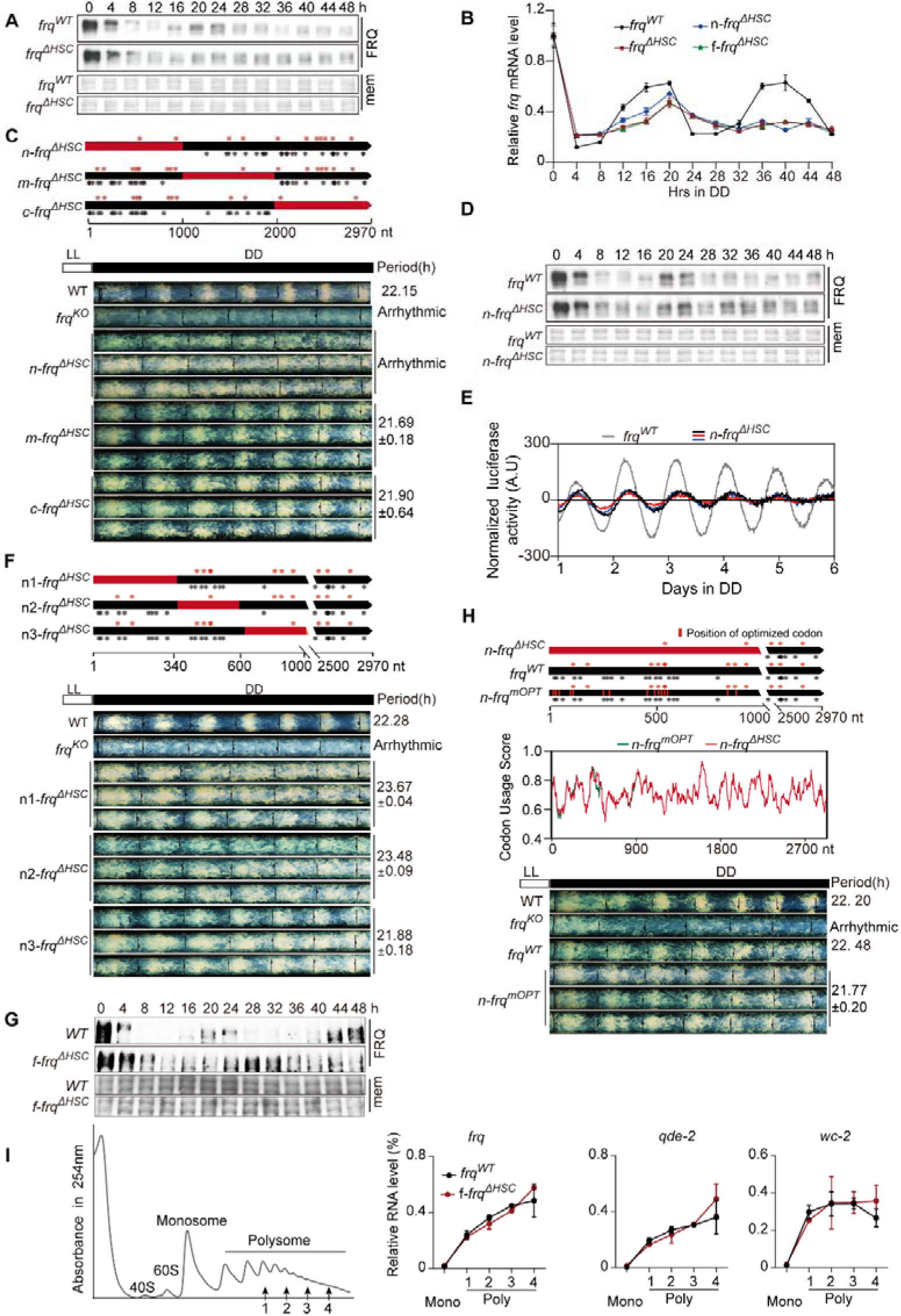
Hidden stop codons (HSCs) in 5’ region of *frq* is important for maintaining circadian rhythm. (A, D and G) Western blot showing the rhythmic expression of the FRQ protein in *frq^WT^*, *frq*^Δ*HSC*^, n-*frq*^Δ*HSC*^ and f-*frq*^Δ*HSC*^ strains upon shifting in the constant dark (DD) conditions. mem, Coomassie Brilliant Blue staining of the PVDF membrane, which serves as a loading control. (B) Reverse transcription qPCR detecting the rhythmic expression of *frq* mRNA in different strains in DD conditions. Strand-specific reverse transcription was performed using primers for *frq* and β*-tubulin*. Transcript levels were normalized by β*-tubulin*. (C, F and H) Top panels, schematic diagrams showing the distributions of HSCs in the *frq* gene of different variants. Black bars indicate *WT* sequence region of *frq*, red bars indicate mutation region. HSCs were marked in red (+1 HSC) and black (-1 HSC) stars. Bottom panels, race tube assay showing the rhythmic conidiations of strains in constant darkness (DD) conditions upon shifting from constant light (LL) conditions, with each line indicating the growth front for every 24 hours. Middle panel (H) is codon usage score of n-*frq^mOPT^,* and 15 codons in this *frq* sequence were slightly optimized. (E) *Luciferase* reporter driven by *frq* promoter showing the rhythm of *frq* promoter activity in *frq^WT^* and n-*frq*^Δ*HSC*^ strains in DD conditions. (I) Polysome profiling for *frq^WT^*and f-*frq*^Δ*HSC*^ strains. Monosome and last 4 polysome fractions were used for RT-qPCR in the right panels. 2 independent replicates were performed for each strain

**Supplementary Figure 2.**
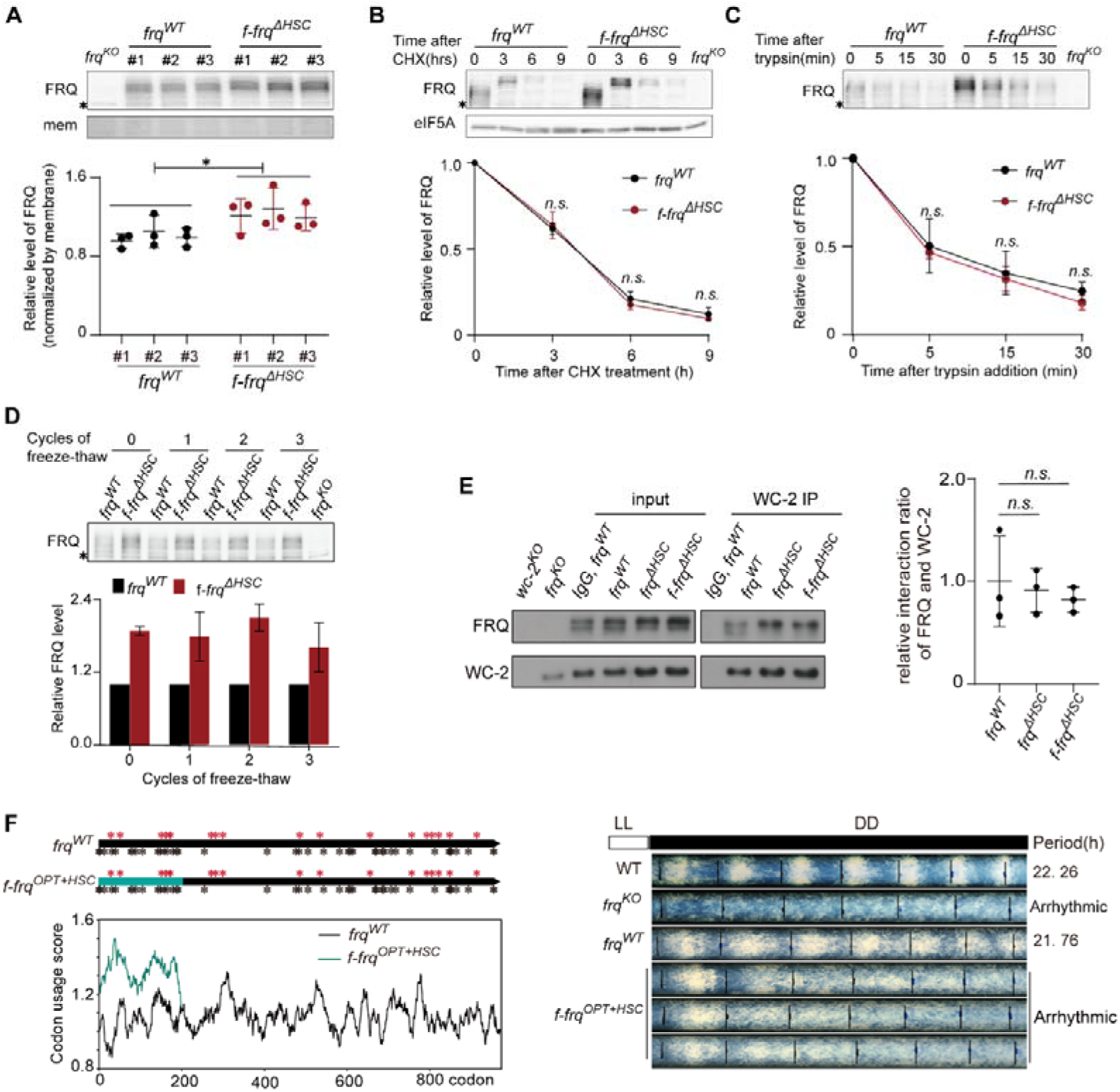
Deleting HSCs in 5’ region of *frq* does not affect the conformation and stability of FRQ. (A) Western blot showing the FRQ protein levels in *frq^WT^* and f-*frq*^Δ*HSC*^ strains in LL conditions. Top panel, western blot results of FRQ, and membrane staining (mem) served as loading control. Bottom panel, densitometric analysis of western blot from 3 independent strains. (B) Degradation of FRQ protein after cycloheximide treatment (10 μg/mL) in LL conditions. Top panel is western blot results, eIF5A served as loading control. Bottom panel, densitometric analysis of western blot result from 3 independent replicates. (C) The sensitivity assay of FRQ protein to trypsin digestion (0.5 μg/mL). Top panel, western blots showing degradation of FRQ protein from *frq^WT^*and f-*frq*^Δ*HSC*^ strains subject to trypsin digestion. Bottom panel is the result of densitometric analysis from 3 biological replicates. (D) FRQ degradation in freeze-thaw cycle assay. Bottom panel is densitometric analysis of western blot (top) from 3 biological replicates. FRQ level of *frq^WT^* strain was served as standard to normalize FRQ level in f-*frq*^Δ*HSC*^ strains in each freeze-thaw cycle. (E) The affinity of FRQ protein to WC-2 in *frq^WT^* and f-*frq*^Δ*HSC*^ strains. Left panel, western blot results from co-IP assay of WC-2 protein. Right panel, densitometric analysis of western blots from 3 independent replicates. The affinity of FRQ to WC-2 was calculated using IP ratio (FRQ/WC-2) divided by the input ratio (FRQ/WC-2). (F) Left top panel, HSCs distribution in *frq^WT^* and f-*frq^OPT+HSC^* sequence. Left bottom panel, codon usage score of *frq^WT^* and f-*frq^OPT+HSC^* sequence. Right panel, race tube assay for f-*frq^OPT+HSC^* strain. Upstream 200 codons in f-*frq^OPT+HSC^* sequence were optimized and maintained their original HSCs. For A-E panels, error bars for standard deviation of three independent biological replicates. *P* values were calculated by unpaired two-tailed *t*-test. *, *P* < 0.05. **, *P* < 0.01. ***, *P* < 0.001. *n.s.*, no significant.

**Supplementary Figure 3.**
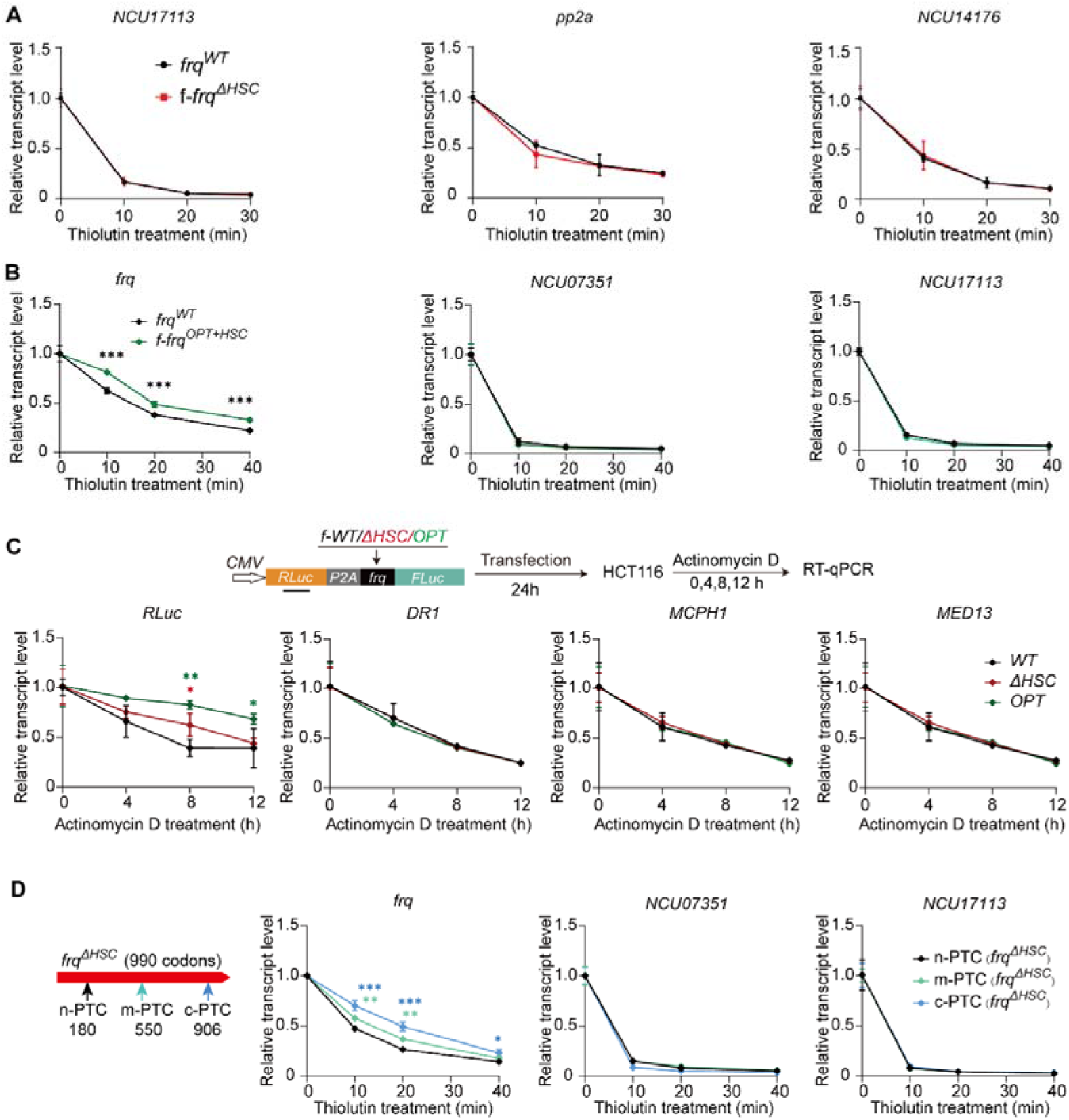
Reverse transcription quantitative PCR (RT-qPCR) detects the mRNA decay. (A) RNA decay assay in *frq^WT^*and f-*frq*^Δ*HSC*^ strains of *N. crassa* treated with thiolutin. The three genes were set as negative internal control of Figure 1I. (B) RNA decay assay in *frq^WT^*and f-*frq^OPT+HSC^* strains of *N. crassa* treated with thiolutin. (C) RNA decay assay for HCT116 cell transfected test plasmid. The transcript, *RLuc*-*frq*-*FLuc* and endogenous genes were quantified using spik-in *Neurospora* β*-tubulin*. (D) RNA decay assay for *frq*^Δ*HSC*^ strains harboring an in-frame stop codon (n-PTC, m-PTC, or c-PTC). Transcripts in A and C were quantified using spike-in human *GAPDH* gene. For A-D, 3 independent replicates were performed. *P* values were calculated by unpaired two-tailed *t*-test (n = 3 independent samples) and One-way ANOVA (C). *, *P* < 0.05. **, *P* < 0.01. ***, *P* < 0.001.

**Supplementary Figure 4.**
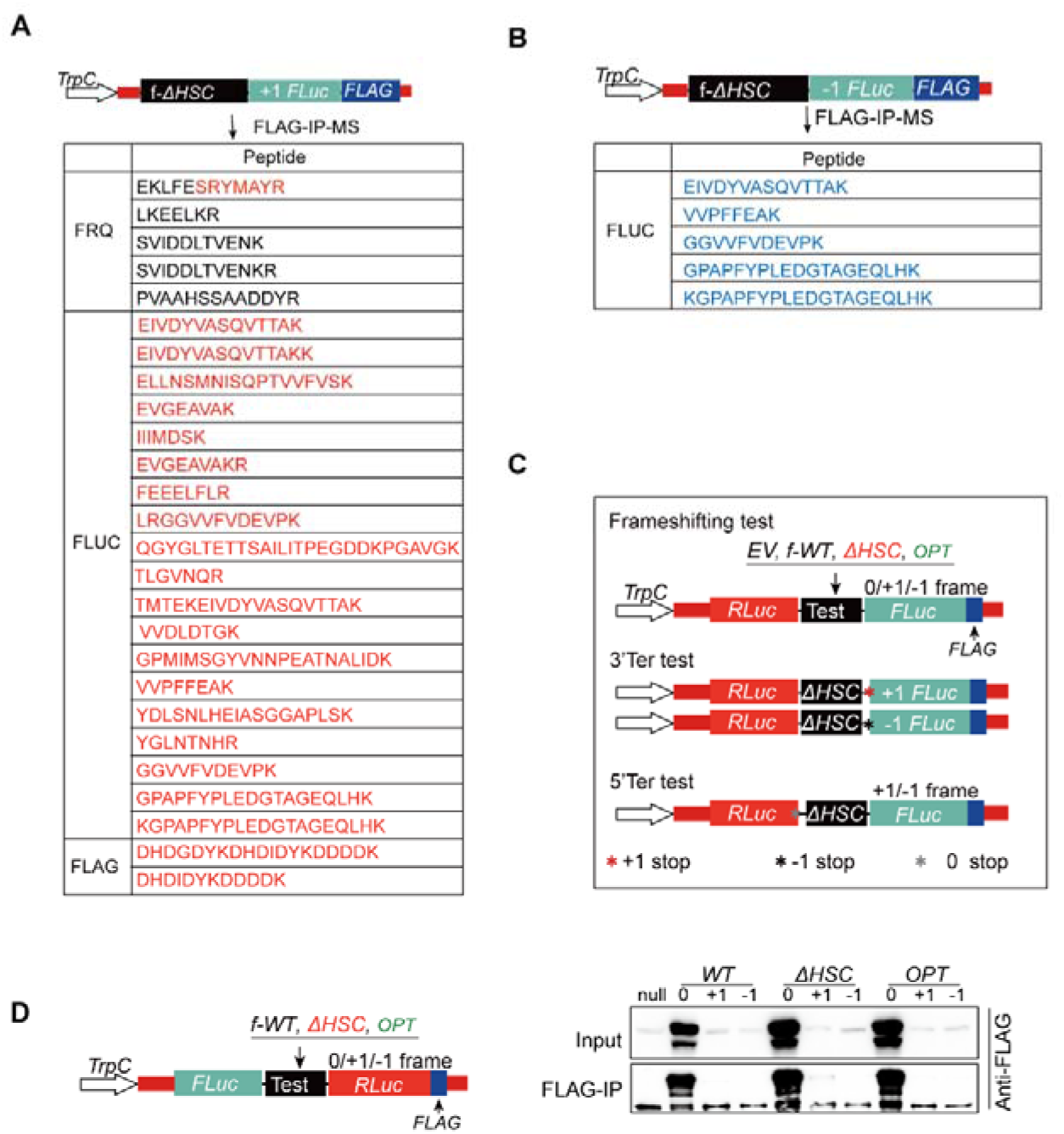
Detection of ribosomal frameshifting in *frq* sequence in *N. crassa in vivo*. (A and B) Mass spectrometry results showing out-of-frame and *trans*-frame peptides detected from strains expressing f-Δ*HSC-FLuc-FLAG*, with *FLuc-FLAG* in +1 and -1 frame, respectively. Peptides sequences marked with black, red and blue are encoded by 0, +1 and -1 frame, respectively. (C) The dual luciferase reporter system for detecting ribosomal frameshifting of *frq* in *N. crassa*. All constructs were driven by *TrpC* promoter. The dual-luciferase reporter has 5’UTR and 3’UTR from *frq* gene, and *frq* sequences (f-*WT,* f*-*Δ*HSC* and f-*OPT*) sequences were inserted between *RLuc* and *FLuc-FLAG*. The *FLuc* sequence was located on 0, +1 or -1 frame. For 3’Ter test structure, a +1 or -1 frame stop codon was inserted after test sequence. For 5’ Ter, 0 frame stop codon was inserted before test sequence. The stars were marked with red (+1 HSC), black (-1 HSC) and gray (in frame stop codon). (D) Left panel, modified dual-luciferase reporter featuring inverted *Rluc* and *Fluc* orientations relative to the construct depicted in supplementary figure 4C. Right panel, FLAG-IP assay to confirm the frameshifting of f-*frq* fragments (*WT*, Δ*HSC* and *OPT*) with the left reporter system.

**Supplementary Figure 5.**
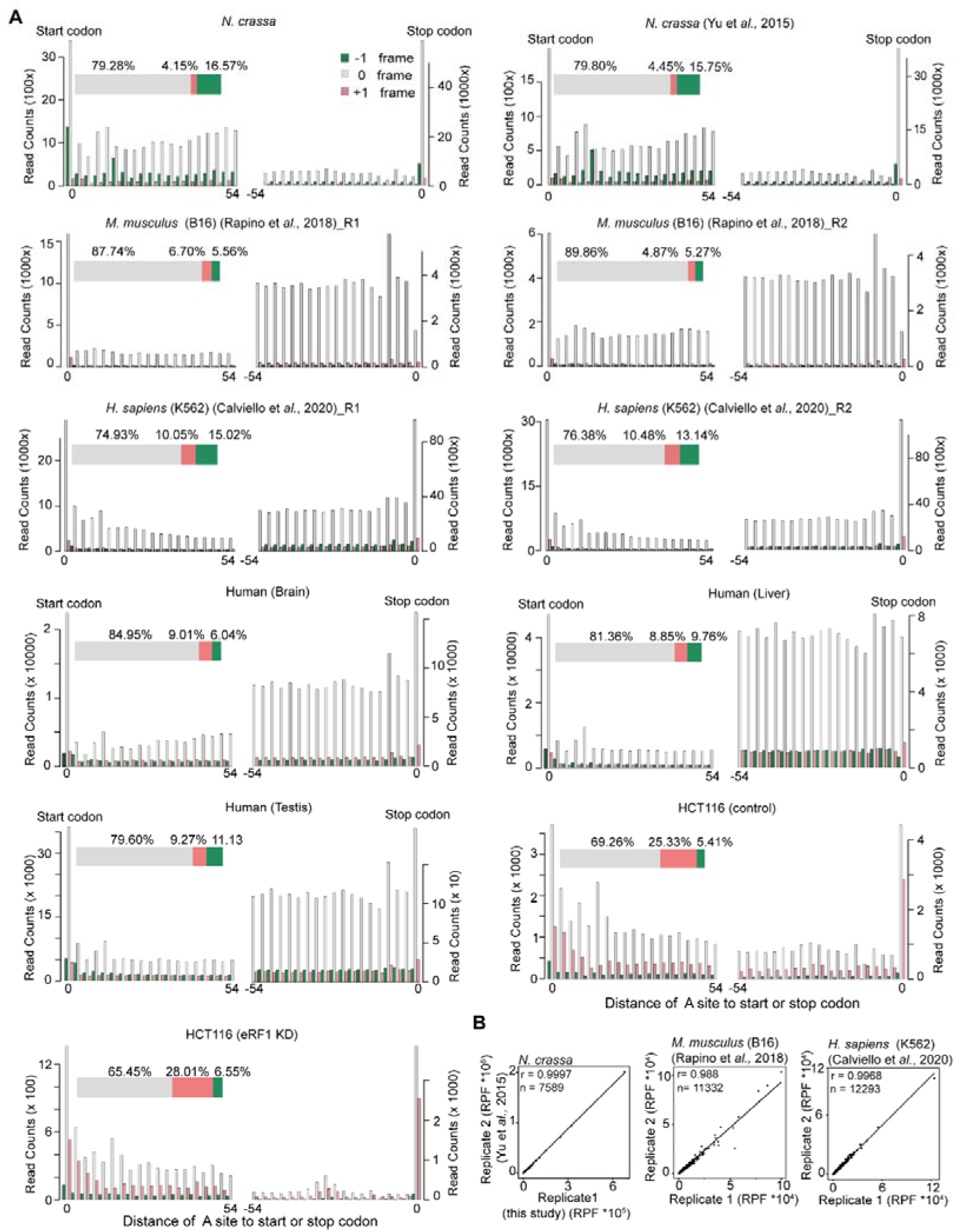
The quality analyses for high-resolution ribosomal profiling sequencing data. (A) Distribution of RFs from the start or stop codon, showing a 3-nucleotide periodicity. RFs are represented in gray (0 frame), pink (+1 frame), and green (-1 frame). Apart from three ribosomal profiling datasets (for *Neurospora* and HCT116) generated in this study, all other Ribo-seq data were obtained from published sources. (B) The repeatability analysis of ribosomal profiling data from different species. For the data of *N. crassa*, replicate 2 was downloaded from published data, and replicate1 was developed in this study.

**Supplementary Figure 6.**
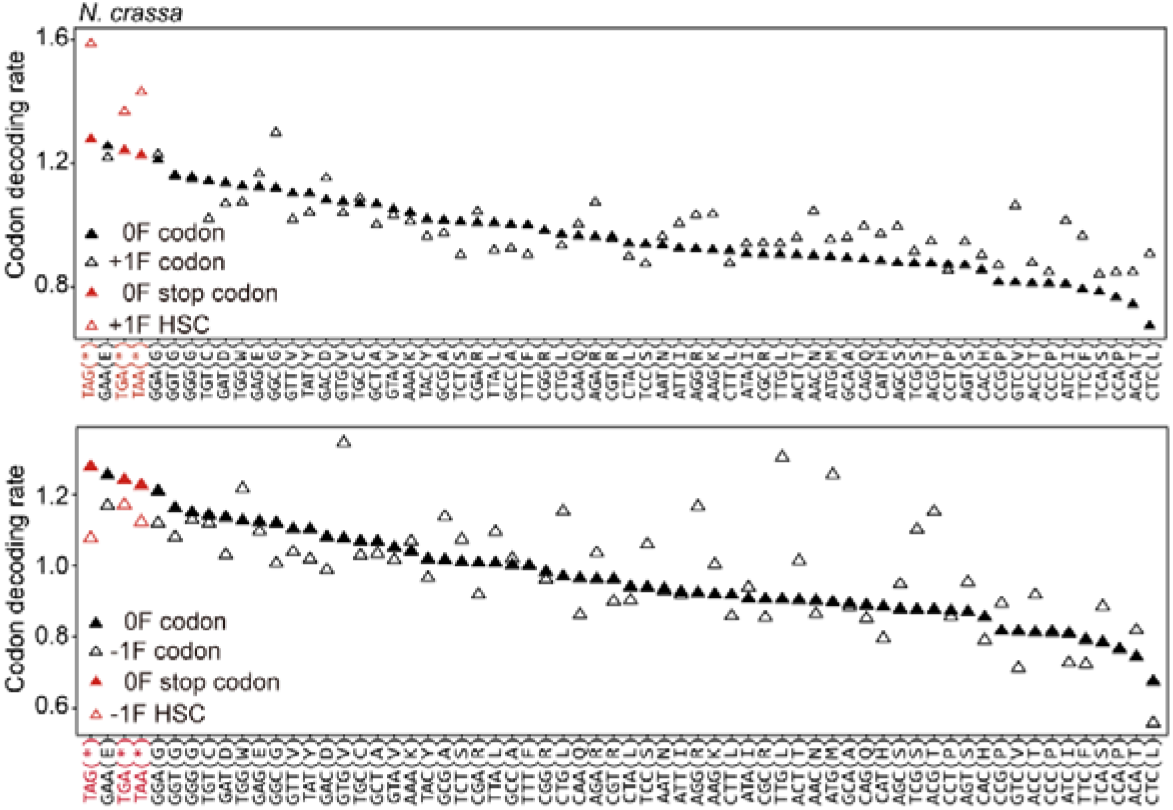
Dot plots of the codon decoding rate (CDR) of 64 codons based on A sites of reads in 0 frame and +1 frame (top panel) or -1 frame (bottom panel). The *N. crass*a ribosomal profiling data was carried out in this study.

**Supplementary Figure 7.**
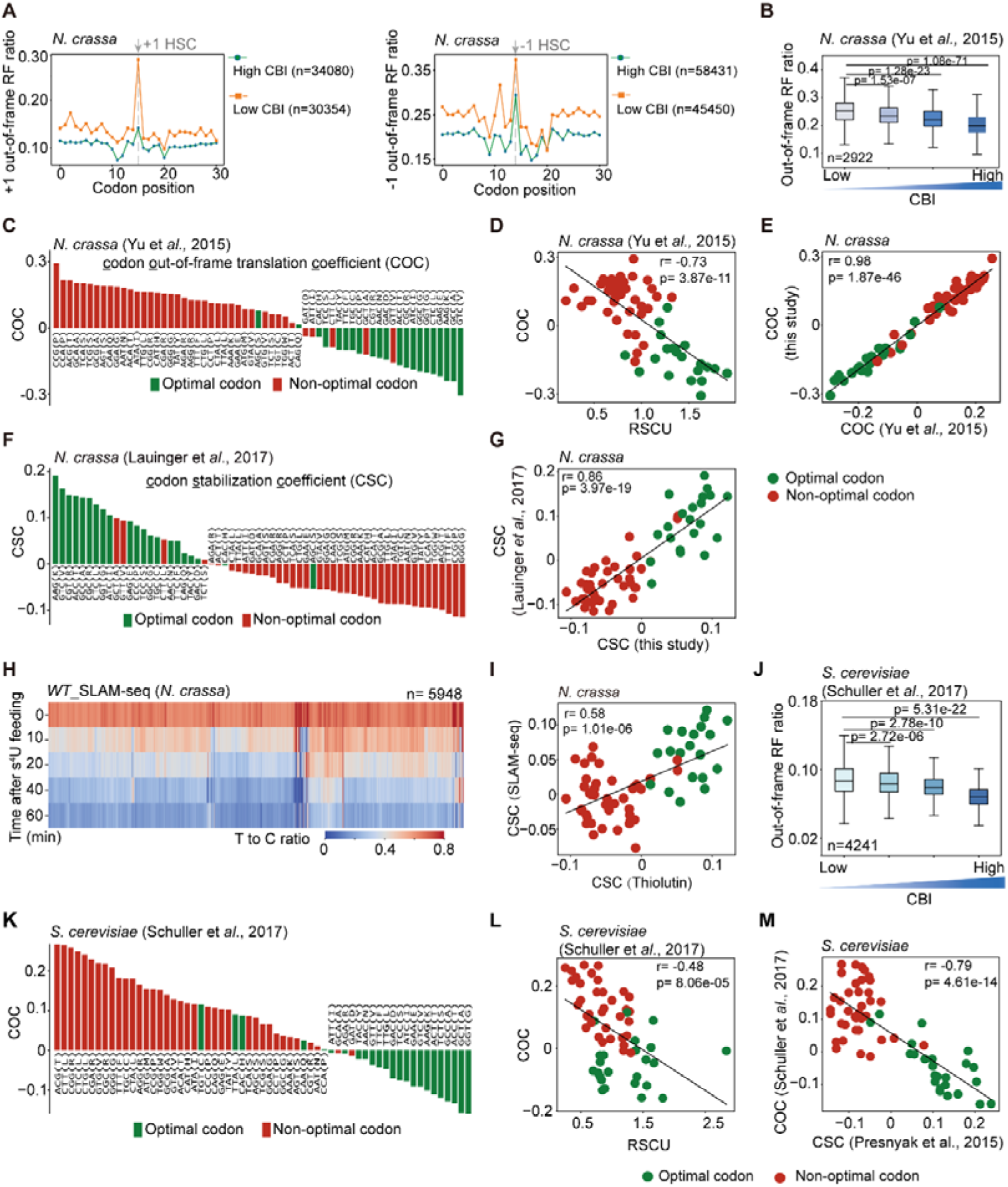
The correlation between mRNA stability and putative out-of-frame translation is conserved in fungi. (A) Line plots showing the distribution of +1 (left panel) or -1 (right panel) out-of-frame RFs flanking respective HSCs based on our Ribo-seq data. Gene fragments were stratified into 4 quantiles based on CBI, with Q1 and Q4 representing low or high CBI group, respectively. Number of fragments harboring HSC were indicated in the bracket. (B) Boxplot showing out-of-frame RF rations of genes stratified into 4 quantiles based on gene codon bias indices and out-of-frame RF ratios from published Ribo-seq data of *N. crassa*. Student t-test was performed (B and J). (C) Bar plot displaying <u>c</u>odon <u>o</u>ut-of-frame translation <u>c</u>oefficient (COC) of each codon based on published Ribo-seq data of *N. crassa*. (D) Scatter plot showing the correlation between COC and RSCU value of 61 sense codons. The COC was obtained from published ribo-seq data of *N. crassa*. (E) The repeatability analysis of COC values across 2 independent ribosomal profiling of *N. crassa*. (F) Bar plot displaying CSC values of 61 sense codons derived from published RNA decay data of *N. crassa* with thiolutin treatment. (G) The repeatability analysis of CSC values across 2 sets RNA decay data from published and developed in study. (H) Heatmap depicting time-resolved T>C conversion rates for *N. crassa* genes across five time points in a SLAM-seq time-course experiment. (I) The repeatability analysis of CSC values across RNA decay data with thiolutin treatment and SLAM-seq. (J) Boxplot showing out-of-frame RF rations of genes stratified into 4 quantiles based on gene CBI values. The out-of-frame RF ratios from published Ribo-seq data of *S. cerevisiae*. (K) Bar plot displaying COC value of each codon derived from published Ribo-seq data of *S. cerevisiae*. (L) Scatter plot showing the correlation between RSCU and COC values in published Ribo-seq data of *S. cerevisiae*. (M) Scatter plot showing the correlation between COC (Ribo-seq data.) and CSC values (RNA decay data from published data) in *S. cerevisiae*.

**Supplementary Figure 8.**
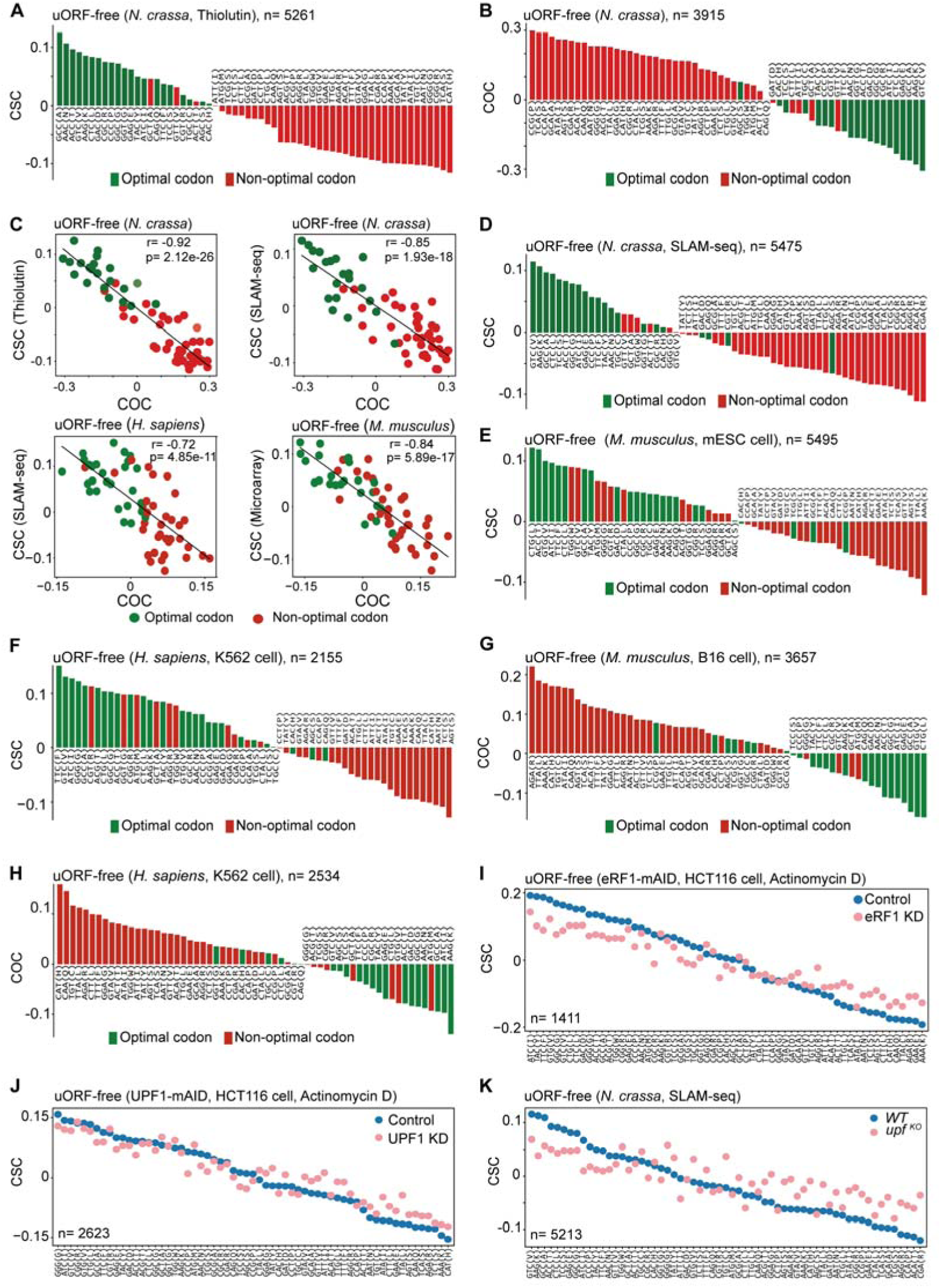
Codon optimality tightly links the mRNA stability and putative out-of-frame translation in uORF-free genes. (A, D, E and F) Bar plots showing the CSC values of 61 codons based on uORF-free genes in RNA decay data. the RNA decay data for mouse mESC and human K562 was downloaded from published data. (B, G and H) Bar plots showing COC values of 61 codons based uORF-free genes in Ribo-seq data. The data for mouse B16 and human K562 was downloaded from published Ribo-seq data. (C) Scatter plots showing the correlation between CSC and COC values which based on uORF-free genes. (I, J and K) Scatter plots of CSC values for 61 codons derived from uORF-free genes in RNA decay profiling. The codons are arranged in descending order based on the CSC values of the control or WT sample. Each point is marked in blue (control or WT) or pink (depletion or knock-out). uORF-containing genes were identified using *Neurospora crassa* Ribo-seq data generated in this study (analyzed via RiboToolkit) or from annotated datasets (*Mus musculus* and *Homo sapiens*).

**Supplementary Figure 9.**
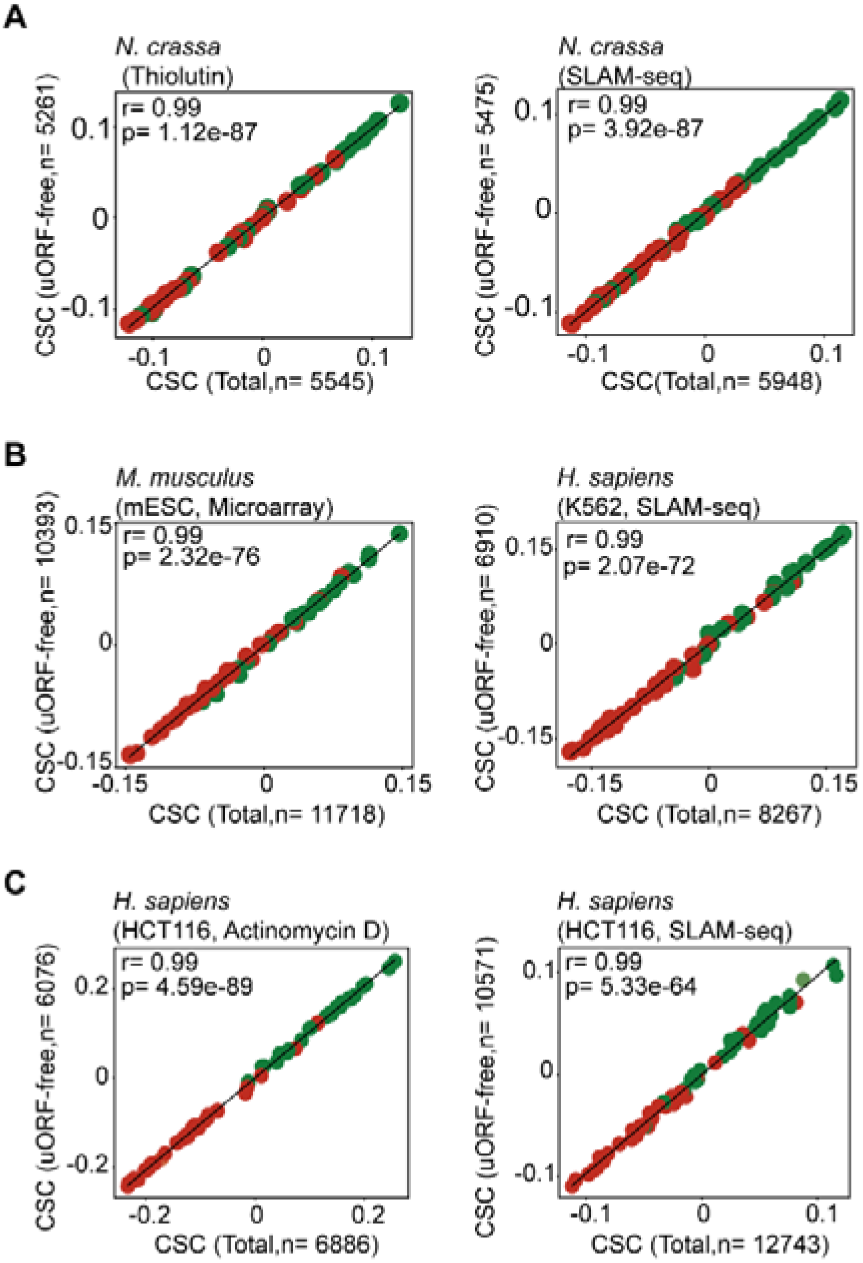
Repeatability between CSC values based on total and uORF-free genes in RNA decay data. uORF-containing genes were identified by analyzing Ribo-seq data using RiboToolkit.. The RNA decay data sets were derived from our data (panel A, *N. crassa* and panel C, HCT116 cell) and published data sets (panel B, mESC and K562).

**Supplementary Figure 10.**
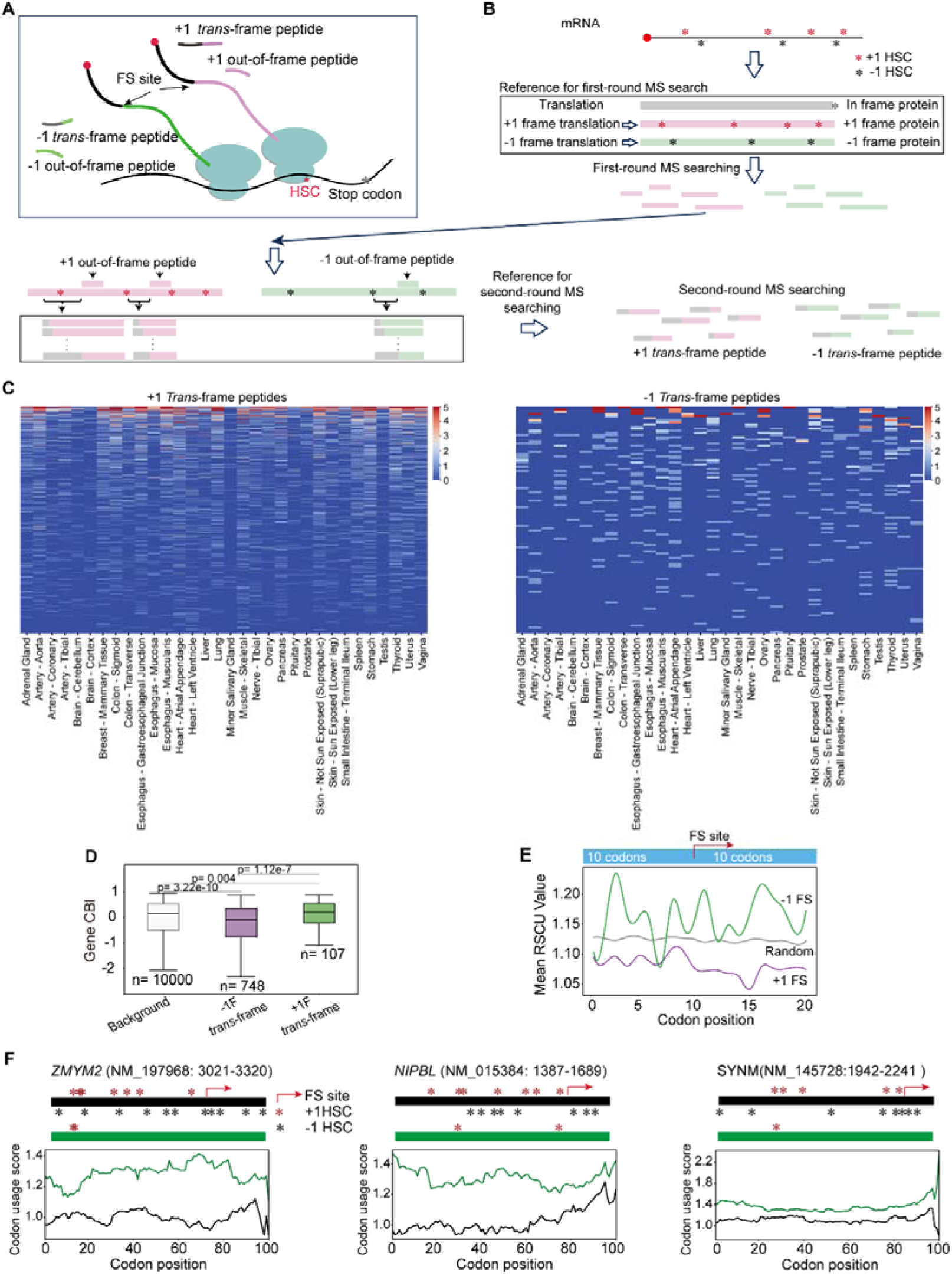
Non-optimal codon usage induces ribosome frameshifting in tissue and cells. (A) Schematic diagram illustrating the definition of out-of-frame and *trans*-frame peptides. (B) Diagram showing the strategy to identify ribosomal frameshifting in human tissues. Firstly, we searched for proteins in the dataset using all annotated proteins from UniProt (UP000005640). The detected proteins were set as background reference. Subsequently, we expanded the reference by incorporating out-of-frame proteins, which are translated from +1 or -1 frame relative to the original ORF (reference was list in Table S2). Using this reference, we performed the first-round searching in mass spectrometry data to identify out-of-frame peptides. By pinpointing the initiation sites of these out-of-frame peptides, we traced the first hidden stop codon (HSC) upstream of the out-of-frame fragment to limit the region of frameshift site. Then, we constructed a *trans*-frame peptides reference with shifting on every codon in this region, and combined with the background reference as a reference for the second-round mass spectrometry detection (reference was list in Table S3). (C) Heatmap of +1 (left panel) or -1 (right panel) *trans*-frame peptides detected across 32 human tissues via mass spectrometry. (D) Boxplot showing the CBI distribution of background and genes with detected *trans*-frame peptides (+1 or -1 frame) in tissue mass spectrometry (E) Line plot showing the mean RSCU value per codon position in 20-codon fragments of random or harboring detected ribosome frameshifting site. (F) HSC distribution (top panels) and codon usage score of ∼300 bp fragments (bottom panels, *WT* or *OPT* variants) from candidates with detected frameshifting sites.

**Supplementary Figure 11.**
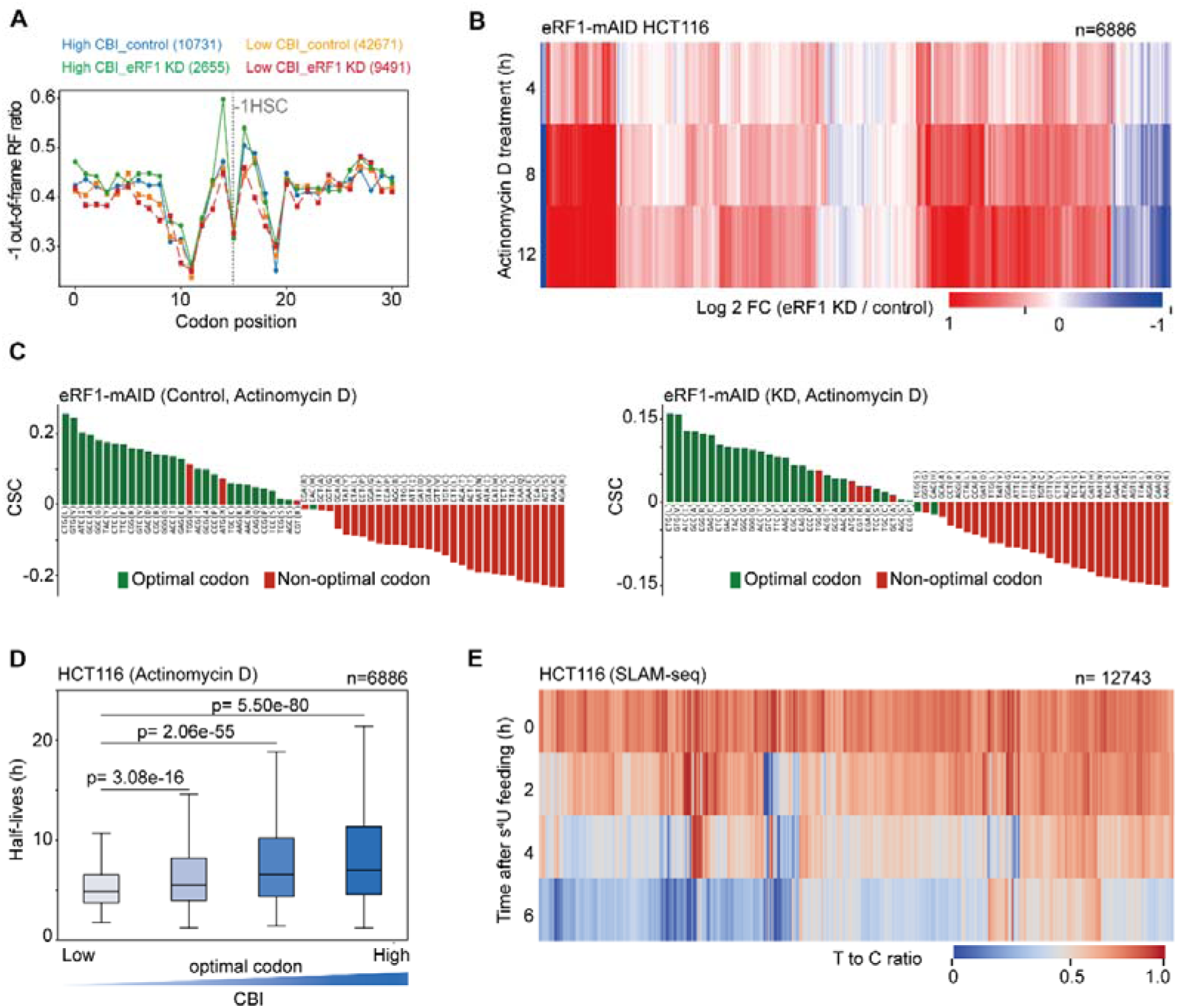
eRF1 involves in frameshifting induced RNA decay. (A) Line plot showing distribution of -1 out-of-frame ribosome footprints around -1 HSC in gene fragments with low or higher CBI, under normal or eRF1 KD condition. Fragments were stratified into 4 quantiles based on CBI, with Q1 and Q4 representing low or high CBI group, respectively. Number of fragments harboring HSC were indicated in the bracket. (B) Heatmap displaying the Log2 fold changes of transcripts upon eRF1 knockdown (KD), compared to the control of each gene in different time points. (C) Bar plots showing CSC values of 61 codons based RNA decay data of control (left panel) or eRF1 depletion (right panel) samples in eRF1-mAID cells. (D) Box plots displaying half-lives of genes stratified into four quantiles based on CBI. The RNA decay data was derived from control sample of eRF1-mAID HCT116 cell treated with actinomycin D. Student t-test was performed. (E) Heatmap depicting time-resolved T>C conversion rates for genes in HCT116 cells across 4 time points in a SLAM-seq time-course experiment.

**Supplementary Figure 12.**
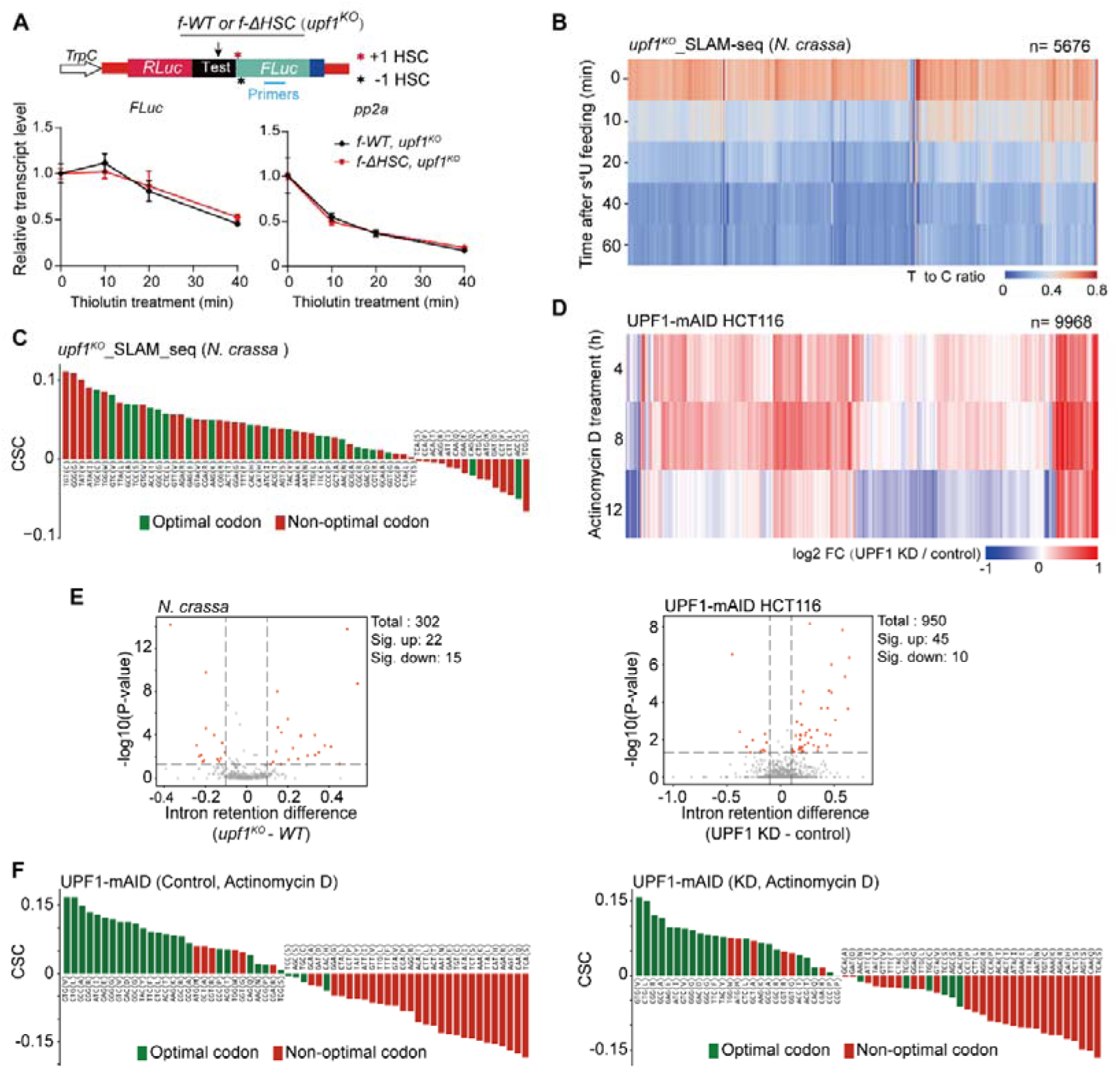
UPF1 involves in regulation of codon usage related mRNA stability. (A) RNA decay assay for *upf1^KO^*strains containing the dual luciferase constructs with f-*WT* or f-Δ*HSC* insertion. The transcript level was quantified by spike-in human *GAPDH* gene. *pp2a* was set as an internal control. (B) Heatmap depicting time-resolved T>C conversion rates for *N. crassa* genes in *upf1^KO^* strain across 5 time points in a SLAM-seq time-course experiment. (C) Bar plot showing CSC values of 61 codons based on SLAM-seq data of *upf1^KO^* strain. (D) Heatmap displaying the Log2 fold changes of transcripts upon UPF1 knockdown (KD), compared to the control of each gene in different time points. (E) Volcano plots showing intron retention difference between WT (or control) and *upf1^KO^*(or UPF1 depletion) in *Neurospora* and HCT116 UPF1-mAID cell line. X-axis: log2-fold change in retention rate; Y-axis: -log10 (adjusted p-value). Red points: significantly retained introns (|log□FC| >1, P value <0.05). Dashed lines: significance thresholds. (F) Bar plots showing CSC values of 61 codons based on RNA decay data of control (left panel) and UPF1 depletion (right panel) samples in UPF1-mAID HCT116 cells. The samples were treated with actinomycin D.

## METHOD DETAILS

### Operation and culture condition of *Neurospora crassa* strains

*N. crassa* Strains harboring different modified *frq* sequences were generated in this study (key resources table). The *frq^KO^* strain was used as host strain for various *his-3*-targeting constructs that contain entire *frq* locus (*frq* promoter, UTR regions and *frq* CDS sequence). The 9716 *his-3^-^*and *upf1^KO^ his-3^-^* strain were used as host strain for various his-3-targeting constructs for dual-luciferase assay.

Mycelia were grown in minimal medium (2% glucose and 1 ×Vogel’s which contains 3 g Na_3_•Citrate•5H_2_O, 5 g KH_2_PO_4_, 2 g NH_4_NO_3_, 0.2 g MgSO_4_•7H_2_O, 0.1 g CaCl_2_•2H_2_O, 250 μg biotin and 1× Trace element in 1 liter). Race tube medium contains 1×vogel’s, 0.1% glucose, 0.17% arginine, 50 ng/mL biotin and 1.5% agar. Strains grow in slants with the medium (1×vogel’s, 2% sucrose and 1.5% agar) in 30°C temperature.

### Human cell culture and transfection

Cells were cultured in DMEM medium supplemented with 10% fetal bovine serum, 1×penicillin-streptomycin and incubated at 37□ with 5% concentration of CO_2_. Cell lines used in this study were list in key resource table.

For transfection, cells were seeded in a 12-well cell-culture dish to get a 70-80% confluency after 16 h prior culturing. Then plasmids were prepared for transiently transfected into cell using a calcium phosphate cell transfection kit. In brief, 1.5 μg plasmid was mixed with 40 μL CaCl_2_ solution. Then DNA-CaCl_2_ mixture was added into 40 μL BBS solution. After incubation at room temperature for 10 min, the DNA-CaCl_2_-BBS mixture was added gently into the cell. After 8 hours, the medium was replaced with fresh pre-warmed medium. For additional 16 h, the cells were ready for protein or RNA extraction.

### Generation of eRF1-AID and UPF1-AID cell lines

For generation of AID plasmid, guide RNA plasmid was developed based on pX330-U6-Chimeric_BB-CBh-hSpCas9. Two donor plasmids containing homologous arm, AID coding sequence, either neomycin or hygromycin resistance gene, were developed based on pMD19-T vector. Plasmids and primers used in this study were list in Table S7 and S9.

HCT116-OsTIR1 cell line ^78^ was used for generation of AID cell line. Cells were cultured in DMEM supplemented with 10% fetal bovine serum at 37 °C in a humidified incubator with 5% CO_2_. Before transfection, cells were seeded in a 6-well plate at 70% coverage. After 24 h culturing, 1.6 µg guide RNA plasmid and 2.4 µg donor plasmids (1.2 µg of each donor plasmids) were transfected into cells using a calcium phosphate cell transfection kit. After 8 h transfection, medium was refreshed. Cells underwent selection with 500 µg/mL G418 and 100 µg/mL hygromycin starting 24 hours post-transfection. After two weeks of continuous selection, viable cells were dispersed and isolated via limiting dilution in 96-well plates to obtain single-cell clones.

For eRF1 or UPF1 depletion, selected clones were incubated with 1 µg/mL doxycycline for 24 h to induce TIR1 expression. Subsequently, 10 µM 5-phenylindole-3-acetic acid (5-Ph-IAA) was added to trigger targeted protein degradation. Protein lysates were then extracted for Western blot analysis.

### Plasmid constructs and *N. crassa* transformation

*pKAJ120* plasmid, containing entire wild-type *frq* gene locus, and *his-3*-targeting sequence, were served as backbone for modified *frq* sequence. Modified *frq* fragments were inserted to parent plasmid by replacing WT *frq* (CDS) sequence. The new constructs were transformed into host strain *frq^KO^*by electroporation and targeted to *his-3* locus.

*Luciferase* reporter construct (*pCSR-1-Pfrq-luc-bar*) were generated by inserting *Pfrq-luc* fragment to *pCSR-1*construct by courtesy of Dr. Yi Liu. The new reporter construct contains *luciferase* gene controlled by *frq* promoter and a *bar* gene for resistance screening. Then the reporter was transformed into WT strain and other *frq* rescued strains by electroporation and targeted to *csr-1* locus. Transformants were selected by basta (glufosinate-ammonium, 200 μg/mL) and Cyclosporin A (5 μg/mL) resistance conferred by *bar* and deletion of *csr-1* gene, respectively.

### Protein extraction and western blot analysis

Protein extraction and western blot analysis for *N. crassa* were performed as previously described. ^86^ In briefly, samples were harvested and grinded in liquid nitrogen, then resuspending sample powder with pre-cool protein extraction buffer (50 mM HEPES [pH 7.4], 137 mM NaCl, 10% glycerol, containing proteinase inhibitor (1 mM PMSF, 1 μg/mL pepstatin, 0.5 mg/mL leupeptin). After incubating on ice for 5 min, samples were centrifuged at 12,000×*g* for 5 min at 4°C.

For extracting protein from cell, cell was lysed with lysis buffer (50 mM Tris-HCl [pH 7.5], 150 mM NaCl, 0.1% Triton-X100, containing proteinase inhibitor, 1 mM PMSF, 1 μg/mL pepstatin, 0.5 mg/mL leupeptin) for 15 minutes at room temperature with gently shaking. The cell lysate was collected and centrifuged at 20,000 × *g* for 5 minutes at 4°C. the supernatant was transferred to new tube for further assay.

The protein concentration was quantified using the Bradford method. The collected protein samples were mixed with protein loading buffer and denatured at 100□ for 10 min. Then denatured proteins were separated on SDS-PAGE gels and transferred to PVDF membranes using wet transfer The membrane with protein blot was incubated in 1 × TBS with 0.3% Tween-20, 0.5 % skim milk and corresponding antibodies.

### RNA isolation and expression level analysis

Total RNA was extracted with Trizol reagent. 1 mL Trizol agent was added into 300 mg sample or 6-well culture dish, then 200 μL chloroform was added to mixture after 5 min rotation. Then the samples were centrifuged for 15 min at 20,000 ×*g* at 4°C. The aqueous phase was transferred to a new tube with 200 μL chloroform. The samples were mixed and centrifuged again. The aqueous phase transferred to a new tube. Then RNA was precipitated with pure ethanol (2 volume of transferred aqueous phase) at 4□ for 2 hours. Samples were centrifuged 20 000 × *g* at 4□ for 15 minutes. The RNA pellet was washed 2 times with 70% ethanol. Air dried RNA pellet was dissolved in RNase free water. RNA concentration was determined with Colibri Spectrometer (TITERTEK BERTHOLD).

Reverse transcription (RT) and real-time quantitative PCR were performed as previously described. ^87^ For detecting *frq* mRNA level, strand-specific RT reaction was performed using HiScript II 1st Strand cDNA Synthesis Kit (Vazyme, R211). 2 pmol RT primers of *frq* and β*-tubulin* were added into reaction. And for detecting other genes, reverse transcription (RT) reaction was performed using HiScript II Q RT SuperMix for qPCR (Vazyme, R223). And qPCR was performed on LightCycler® 480 Instrument (Roche). qPCR primers used in this study were listed in Table S9.

### Luciferase reporter assay for circadian rhythm

Strains harboring *Pfrq-luc* reporter were monitored by LumiCycle (Actimetrics) to observe the rhythmicity of *luciferase* reporter with a published protocol. ^20^ The AFV (autoclaved FGS-Vogel’s) medium contained 1×FGS (0.05% fructose, 0.05% glucose, 2% sorbose), 1×vogel’s medium, 50 μg/L biotin, 1.8% agar and 50 μM firefly luciferin. Firefly luciferin was added to medium after autoclaving. Conidial suspensions were spotted onto AFV medium and incubated overnight (30°C, light). Cultures were then transferred to constant darkness (25°C) in LumiCycle instruments for continuous bioluminescence recording. Raw luminescence data were baseline-normalized (24h moving average subtraction) using LumiCycle Analysis software (Actimetrics). Notably, within our experimental conditions, the luciferase signals displayed variability during the initial day within the LumiCycle apparatus, but stabilized thereafter. Consequently, the recorded results were documented after one day under constant dark conditions.

### Race tube assay

Race tube assay was followed with the published protocol. ^20^ Race tube is a glass tube with 40 cm length. Both ends of tube are upswept about 60° and clogged with cotton balls. 13 mL race tube medium were added into race tube. Inoculating mycelia onto medium at one end and allowing mycelia to propagate towards the opposite side. Then cultures were placed under constant light (LL) condition at 25°C for 24 h, subsequently shifted to constant darkness (DD) at 25°C. The growth front of mycelium was recorded at every 24-h period upon shifting in to DD condition. Circadian period (τ) was calculated using τ (h) = [Distance between conidiation bands (mm)] / [growth length(mm)] ⅹ 24 h.

### Detecting rhythmicity of *frq* mRNA and FRQ protein

The method for detecting *N. crassa circadian* in liquid culture was followed as previous described. ^88^ Spreading conidia on 2% glucose medium in culture dish to get a mycelia mat, then mycelia disks were excised from the mat. One such disk was then transferred to 50 mL 2% glucose medium and cultures were grown under constant light with 100 rpm shaking at 25°C. The cultures were transferred to darkness, and underwent a period of growth for some hours before being harvested. Harvesting was performed at 4-hour intervals across a 48-hour dark incubation period (13 timepoints: 0–48 hours). Each timepoint represents an independent biological replicate cultured under identical conditions. Crucially, all cultures received identical growth duration from initial culture to harvesting.

For FRQ protein rhythm, extracting total protein and diluted to same concentration for protein extracts, then equal amount of protein (40 μg) was used for western blot. For rhythm of *frq* mRNA, the total RNA was extracted using Trizol reagent. Dissolving RNA pellet with DEPC treated water and diluting to same concentration. Strand-specific RT reaction was performed using equal amount of RNA.The relative level of *frq* mRNA was quantified by internal control β*-tubulin*. The sample of 0-h time point serves as the standard for normalization.

### Protein stability assay

FRQ protein stability assay was performed as describe. ^20^ Mycelial disks were cultured in 50 mL of 2% glucose medium under constant light for 24 h prior to harvesting or treatment. For cycloheximide (CHX) inhibition assay, Cultures were treated with CHX (10 μg/mL final concentration). Samples were harvested at 0, 3, 6, and 9 h post-treatment. FRQ protein levels were quantified by Western blot and normalized to eIF5A, with the 0-h timepoint serving as the baseline. For detecting FRQ stability using freeze-thaw assay, samples were harvested and grinded to powder in liquid nitrogen. 200 μL protein extraction buffer (without proteinase inhibitor) was added for extracting the protein. The protein was subsequently diluted to a concentration of 2.5 μg/μL. During each freeze-thaw cycle, 200 μL protein of each sample was quickly frozen in liquid nitrogen, then thawed at 37□ for 15 min. 20 μL sample was taken out in a new tube from each freeze-thaw cycle for western blot analysis. For trypsin-sensitivity assay, protein was extracted using protein extraction buffer (without proteinase inhibitor) as previously description. The protein extracts were then diluted to a concentration of 2.5 μg/μL. Subsequently, 100 μL extract was treated with trypsin (final concentration 0.5 μg/μL) at 25□. Aliquots (20 μL) were collected at 0, 5, 15, and 30 min, immediately mixed with loading buffer to terminate digestion, and analyzed by Western blot. FRQ levels at 0 min served as the normalization baseline.

### RNA synthesis *in vitro*

For *RNA synthesis* in vitro, two kinds of reporter assays were used (Table S8): the *renilla luciferase* and *frq-firefly luciferase*. These reporter genes are fused to *psychrophilic phage VSW-3* promoter. ^89^ A 34-nucleotide poly-A tail was appended to the 3’ UTR of the reporter construct via PCR amplification with poly-dT-containing primers. Purified PCR product was subsequently used as templates for *in vitro* transcription with *VSW-3* transcription kit. Following the protocol from manufacturer, 400 ng DNA template was added into 10 μL reaction system. The reaction was incubated at 25□ for 12 hours. Transcription product was extracted with acid phenol-chloroform extraction (volume ratio of 1:1). Then RNA was precipitated with 2 ⅹ volume of pure ethanol, then RNA pellet was washed with 70% ethanol and dissolved in DEPC water. 5 μg purified RNA was capped using the Vaccinia Capping System at 37□ for 1 h. Repeated phenol:chloroform extraction and ethanol precipitation for capped RNA purification. The integrity and quantity of synthesized transcripts was evaluated by denature agarose gel electrophoresis.

### *In vitro* translation

The preparation of *N. crassa* cell-free translation extracts were performed as previous protocol. ^90^ In briefly, *N. crassa* conidia were inoculated into 1 L Vogel’s sucrose medium (1× Vogel’s salts, 2% sucrose) at 1×10□ conidia/mL and incubated at 32°C with 200 rpm shaking under constant light for 6 h. Germinating conidia were harvested by vacuum filtration, resuspended in 1 L pre-warmed Vogel’s medium, and incubated for an additional hour at 32°C with orbital shaking. After re-harvesting, mycelia were rinsed with ice-cold Buffer A (30 mM HEPES-KOH [pH 7.6], 100 mM KOAc [pH 7.0], 3 mM Mg (OAc)_2_ [pH 7.0], 2mM DTT), weighed, and flash-frozen in liquid nitrogen. Frozen tissue was pulverized in a pre-chilled mortar while slowly adding Buffer A (1 mL/g tissue), with the resulting powder transferred to chilled centrifuge tubes and thawed on an ice-water slurry for 1 h. Following centrifugation at 30,600 × g (4°C, 15 min), The supernatant (avoiding the pellet and fatty upper layer) was transferred to new pre-chilled tubes and subjected to a second identical centrifugation step. Clarified extracts were desalted using Zeba™ spin columns (10 mL, Thermo #89882) per manufacturer instructions, supplemented with protease inhibitors (final concentrations: 1.25 mg/mL AEBSF, 0.25 mg/mL each of Pepstatin A, Antipain, Chymostatin, and Leupeptin, plus 0.05 mg/mL Elastatinal), aliquoted (100 μL), flash-frozen in liquid nitrogen, and stored at -80°C.

The *in vitro* translation assay was carried out with the previously described protocol. ^90^ 10 μL reaction mix was prepared, consisting of 5 μL cell-free extract, 1 μL 10×energy mix (200 mM HEPES-KOH [pH 7.6], 10 mM ATP, 1 mM GTP, 200 mM phosphocreatine, 20 mM DTT), 0.12 μL creatine phosphokinase solution (10 mM HEPES-KOH [pH 7.6], 1 mM DTT, 5 U/μL creatine phosphokinase [sigma, C375], 50% glycerol), 0.5 μL 1 M KOAc, 0.2 μL 50 mM Mg(OAc)_2_, 0.1 μL 1 mM amino acid mix (promega, L4461), 0.25 μL RiboLock RNase inhibitor (Thermos Fisher, EO0384), 1 ng control mRNA (*renilla luciferase*) and 10 ng test mRNA (*frq-firefly luciferase*). The reaction mixture was then incubated at 26°C for 30 min. Then, 2.5 μL 5×passive lysis buffer (Promega, E1941) was added to stop translation.

Dual-Luciferase Reporter Assay System (Promega, E1910) was used to measuring the *Renilla* Luciferase (RLuc) and Firefly Luciferase (FLuc) activity of *in vitro* translation product. 2 μL reaction mixture was used to measure the luciferase activity using GloMax 20/20 Luminometer (Promega, E5311). 10 μL LARII and 10 μL stop buffer were used in each reaction. The out-of-frame translation ratio is calculated with the following functions:

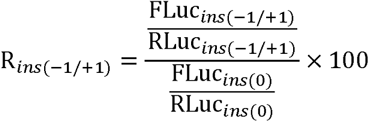

Here, FLuc_ins(-1/+1)_ and RLuc_ins(-1/+1)_ refers to the firefly or Renilla luciferase activities from the constructs with *FLuc* in alternative frames, whereas FLuc_ins(0)_ and RLuc_ins(0)_ refer to firefly or Renilla luciferase activities from the construct with *FLuc* in the 0 frame.

### Dual-luciferase assay *in vivo* of *N. crassa*

Transcript constructs were engineered to monitor ribosomal frameshifting within the *frq* fragment *in vivo* (as depicted in Table S9). In the first reporter systerm, *frq*1-600 bp fragment was inserted between upstream *renilla luciferase* (*Rluc*, codon optimization) and downstream *firefly luciferase* (*Fluc*) positioned in either the 0, +1, or -1 reading frame. For the second systerm, swapping the position of *Rluc* and *Fluc*. All transcripts featured 5’ and 3’ UTRs derived from the native *frq* gene and were driven by the constitutive *TrpC* promoter from *Aspergillus nidulans*. In this assay, three distinct variations of the f-*frq* fragment were utilized: f-*WT,* f-Δ*HSC* (deletion of all HSCs), and f-*OPT* (codon optimization without HSCs). Additionally, an *EV* construct lacking the test sequence was designed as a blank control. In order to excluding internal initiation and aberrant splicing, two distinct constructs have been devised: the 5’ termination control (5’Ter) and the 3’ termination control (3’Ter). For 5’ Ter construct, a stop codon in 0 frame was inserted before insert sequence. And for 3’ Ter construct, a +1 or -1 frame stop codon was inserted in front of +1 or -1 frame *FLuc*, respectively. All of these constructs were subsequently transformed into the 9716 *his-3^-^* strain using electroporation and integrated within the *his-3* locus. The expression plasmids were listed Table S8.

For measuring the luciferase activity in *N. crassa*, *N. crassa* conidia are inoculated into 40 mL 2% glucose medium, with a conidia concentration of 5 μg/mL. The cultures were then incubated at 25°C with 150 rpm under constant light for 36 h. Then, the tissue was harvested, ground into powder in liquid nitrogen, and the total protein was extracted by protein extraction buffer. The luciferase activity was measured by Dual-Luciferase Reporter Assay System (Promega). Frameshifting ratios were evaluated identically to the in vitro assay, using the Relative Luciferase Activity (RLA) of the 0-frame construct as the normalization reference for out-of-frame RLA values.

### Dual-luciferase assay in human cells

A dual-luciferase system was performed to detect the ribosomal frameshifting of test sequences in human cell. ^43^ The pCDNA3.1 plasmid was performed as backbone. Three distinct structures (RF system, P2A system and mP2A system) were performed for frameshifting assay (Table S8). All structures were driven by CMV promoter, and test sequence is inserted between *Rluc* (upstream) and *Fluc* (downstream, placed on 0, +1 or -1 frame) An *EV* construct lacking the insertion sequence was designed as a blank control. Furthermore, the 5’Ter and 3’Ter constructs, identical to those used in the *N. crassa* assay, were also included.

Following transient transfection, cell lysis and luciferase activity measurements were performed using the Dual-Luciferase® Reporter Assay System according to the manufacturer’s protocol. Briefly, cells in 12-well were lysed with 100 μL 1×passive lysis buffer and incubated at room temperature for 15 min. The lysate was centrifuged at 20,000 × g for 2 min at 4°C. The supernatant was used for subsequent luciferase activity measurements. Luciferase activity quantification and ribosomal frameshifting ratio calculations followed the same methodology as used in the *N. crassa*.

### Co-IP assay in *N. crassa*

Samples were suspended in cold protein extraction buffer with protease inhibitor. 2 mg total protein was prepared in 1 mL volume. 50 μL protein was transferred into a new tube as input. Protein G Dynabeads (Thermo Fisher, #10003D) were washed twice with 500 μL of cold protein extraction buffer. After removing the buffer, the beads were resuspended in 200 μL of protein extraction buffer. Next, 2.5 μL of anti-WC-2 antibodies (this study) or control rabbit IgG was added to 12 μL (original beads volume) of the washed beads, then the beads were rotated at room temperature for 45 min to conjugate antibody on beads. Subsequently, buffer was removed, and the beads were resuspended in 50 μL of protein extraction buffer. The prepared protein was transferred to the antibody-bound beads and rotated at 4°C for 3 h. The beads were washed five times with protein extraction buffer and resuspended in 50 μL of 1×SDS loading buffer.

### Mass spectrometry analysis

For FLAG pull-down protein samples were resolved on SDS-PAGE gels and stained with Coomassie Brilliant Blue R-250. Bands corresponding to +1 and -1 out-of-frame proteins were excised for mass spectrometric analysis. Raw MS data for f-Δ*HSC-FLuc-FLAG* samples are deposited in the ProteomeXchange Consortium (PXD044949). Out-of-frame peptides were identified using MaxQuant (v1.6.5.0) against a custom database containing all possible frameshifted variants of f-Δ*HSC-FLuc-FLAG* generated by +1/-1 ribosomal shifting at each codon position. Search analysis using MaxQuant ^91^ built-in default parameters with methodology as we recently described. ^43^

To detect genome-wide ribosomal frameshifting in human tissues, we analyzed mass spectrometry data from 32 normal tissues across 14 healthy donors sourced from the GTEx project (ProteomeXchange: PXD016999 ^73^). Our tiered search strategy first identified expressed proteins against the UniProt human reference proteome (UP000005640) to establish baseline expression profiles. We then constructed an expanded database incorporating theoretical out-of-frame proteins translated via +1/-1 ribosomal frameshifts relative to annotated start codons (Table S2). Initial searches against this composite database identified candidate out-of-frame peptides, whose initiation sites were mapped to pinpoint the first upstream in-frame hidden stop codon (HSC) – thereby defining putative frameshift boundaries. Within these HSC-delimited regions, we generated comprehensive trans-frame peptides covering all possible codon positions (Table S3) and performed a refined validation search against this targeted reference to confirm frameshift events.

Protein identification was performed using MaxQuant (v1.6.5.0) with parameters consistent with our established methodology. ^43^ Each TMT 10-plex experiment analyzed 8 tissue samples alongside 2 reference channels, with 28 independent mass spectrometry acquisitions conducted in duplicate. Database searches incorporated fixed cysteine carbamidomethylation and variable modifications including TMT 10-plex labeling (peptide N-termini and lysine residues), protein N-terminal acetylation, and methionine oxidation. Following peptide-spectrum matching, tissue-specific intensities were normalized against the mean intensity of reference channels. Peptides exhibiting tissue intensity exceeding the reference mean were classified as positive identifications. All detected out-of-frame and trans-frame peptides are comprehensively cataloged in Table S6.

For analyzing codon optimality in frameshifting site, we extracted the central codon plus 10 flanking codons upstream/downstream (21-codon window) in all detected genes with trans-frame peptide. Genome-wide background controls were generated by randomly sampling 10,000 20-codon fragments from all protein-coding transcripts. Codon usage score (average value of the RSCU) or optimal codon proportion of selected gene fragments were calculated.

### Ribosomal profiling

For *N. crassa* ribosomal profiling, the assay followed the published procedure. ^22^ In brief, fresh *N. crassa* conidia (2 × 10□ conidia/mL) were germinated in 200 mL medium (2% glucose) at 25°C with orbital shaking (200 rpm) under constant light for 12 hours. The cultures were harvested via vacuum filtration, and the mycelium pads were ground into powder in liquid nitrogen. Ice-cold 1× yeast polysome buffer with 0.5 mg/mL cycloheximide was added into the tissue powder in equal volume. The mixture was gently mixed and incubated on ice for 5 minutes. The tissue lysates were treated with ARTseq nuclease and then subjected to ultracentrifugation on a sucrose cushion following the ARTseq protocol. The pelleted monosomes were filtered through an Amicon-100K micro-concentrator (Millipore) for 10 minutes at 10,000 × *g*. Then, 500 μL of ribosome release buffer (20 mM Tris-HCl [pH 7.0], 2 mM EDTA, 40 U/mL SUPERase•In) was added to the column, and each sample was incubated on ice for 10 minutes before centrifuging again. The flow-through fraction was collected for RNA purification. The extracted RNA was precipitated with ethanol and glycogen. mRNA footprints of 28-30 nucleotides were purified using urea-PAGE. Library generation was performed according to the ARTseq protocol, and biotinylated rRNA probes were used to reduce rRNA contamination during the library preparation.

For ribosome profiling with human cell line, the assay followed the published protocol. ^44^ 2×10□ HCT116 cells were lysed in 300 µL ice-cold lysis buffer (20 mM Tris-HCl, 150 mM NaCl, 5 mM MgCl□, 1 mM DTT, 100 μg/mL cycloheximide, 0.5% Triton X-100, 2 U/mL DNase I, 1× protease inhibitor cocktail). Lysates were centrifuged at 10,000 × *g* for 10 min at 4°C. Supernatants were treated with RNase I (1 U per OD□□□ unit) for 45 min at 25°C. Total RNA was extracted using TRIzol reagent, precipitated with glycogen, and washed with 80% ethanol. Ribosome-protected fragments (28-34 nt) were resolved on 16% TBE-urea PAGE, excised, and eluted in RNA Gel Extraction Buffer (300 mM NaOAc, 1mM EDTA, 0.25% SDS). RNA was precipitated with 1× volume isopropanol at -20°C overnight. For library preparation, 100-200 ng RNA fragments were adenylated using poly(A) polymerase and PNK (37°C, 30 min). Pre-adenylated 5’ adapter were ligated to ribosome-protected fragments with T4 RNA Ligase 2 truncated KQ (25°C, 2 h). Then purifying the ligation product with Streptavidin magnetic bead.

And template-switching primer was used in reverse transcription with SuperScript III reverse (50°C, 30 min), following the library amplification.

### Ribosome profiling analysis

Ribosome profiling data for *Neurospora crassa* (assembly NC12), *Mus musculus* (GRCm38), and *Homo sapiens* (GRCh38) were processed according to established pipelines. ^22^ Raw sequencing reads underwent adapter trimming and size selection (retaining 25-40 nt fragments), followed by rRNA depletion. Filtered reads were aligned using Bowtie to custom genomic references containing the longest transcript per gene plus 100-nt flanking regions at both ends. By following the methodology described by Ingolia *et al.* ^92^, ribosome-protected footprints (RPFs) were stratified by length, with A-site codon assignment to nucleotides 16-18 from the 5’ end. Subsequently, the groups of RPFs with a high ratio of in-frame codons (e.g., >70% for *N. crassa*, >70% for *H. sapiens*, >80% for *M. musculus*) are selected for further analysis. These groups include RPFs in the 0 frame, as well as the +1 and -1 frames.

Codon decoding rates (CDR) for A-site positions in 0, +1, and -1 reading frames were calculated by normalizing out-of-frame ribosomal footprint (RPF) signals using the average signal of corresponding frame-specific RPFs per gene. Normalized signals for each codon across all genes were summed and averaged based on codon frequency. To minimize terminal artifacts, we excluded out-of-frame RPF signals within 30 nt of the coding sequence (CDS) 3’ end that exceeded the gene mean by >50 normalized units. Genes with fewer than 10 high-quality out-of-frame reads were excluded from analysis. In-frame (0-frame) signals underwent identical processing, but preserved terminal regions (30 nt at both 5’ and 3’ ends) to capture natural termination effects.

To analyze the out-frame ratio for each gene, we calculated the proportion of out-of-frame signals relative to total ribosome-protected footprints (RPFs). Genes with fewer than 200 RPFs across all reading frames (0, +1, and -1) were excluded from subsequent analysis.

### RNA decay assay in *N. crassa*

RNA decay assay in *N. crassa* was performed as previously described. ^57^ Mycelium disks were cut from *N. crassa* mycelia mat, and transferred to 200 mL 2% glucose medium. The cultures were then subjected to shaking at 100 rpm under constant light at 25°C for 24 h. Subsequently, cultures were transferred to 100 mL fresh 2% glucose medium supplemented with 12 μg/mL thiolutin, maintaining identical incubation conditions. Samples were harvested at 0, 10, 20, 30, and 40 minutes post-thiolutin addition, with three biological replicates per time point. To determine RNA degradation profile, 600 ng *N. crassa* RNA and 100 ng HEK293T total RNA (spike-in) were input in RT reaction. For *frq* RNA, gene-specific primers of *frq* gene and human *GAPDH* gene were employed in RT reaction. For other genes, random RT primers were employed in RT reaction. The relative levels of transcripts were subsequently quantified with human *GAPDH* gene.

### RNA decay assay in human cell

RNA decay assay in human cell was performed as previously described. ^5^ The HCT116-eRF1-AID or HCT116-UPF1-AID cell lines were induced by 1 µg/ml of doxycycline for 24 h, and were added or not added (as the control) 10 µM 5-Ph-IAA in medium. Then transferring cultures to fresh medium supplemented with 10 μg/mL actinomycin D (HY-17559, MCE), 10 µM 5-Ph-IAA and 1 µg/ml of doxycycline. Samples were harvested at time points of 0, 4, 8 and 12 hours after actinomycin D treatment. Three independent replicates were collected at each time point. To determine decay rate of genes, 500 ng RNA and 50 ng *N. crassa* RNA were input in RT reaction. Random RT primers were employed in RT reaction. The relative levels of transcripts were subsequently quantified using spike-in *N. crassa* β*-tubulin*.

To assess mRNA stability of test gene fragments under eRF1 knockdown conditions, 1.5 μg plasmid was transfected into HCT116 eRF1-AID cells in 12-well culture dish using polyethylenimine (PEI) for 24 hours. Subsequently, 5-Ph-IAA was added to the medium to deplete eRF1 for 12 hours. Total RNA was extracted with TRIzol reagent, and target mRNA transcript levels were normalized to the *RLuc* gene driven by the *SV40* promoter.

### SLAM-seq assay

The SLAM-seq assay was performed in *N. crassa* and HCT116 cells as previously described. ^67^ For 4-thiouridine (s^4^U) labeling in *N. crassa*, mycelial disks were cultured in 200 mL of 2% glucose medium under light for 24 hours. Subsequently, 100 μM s^4^U was added to the medium for a 2-hour pulse labeling. After labeling, the disks were washed three times with fresh medium using vacuum filtration, then transferred to fresh medium supplemented with 10 mM uridine to terminate s^4^U incorporation. Cultures were harvested at 0, 10, 20, 40, and 60 minutes after transfer. For labeling in HCT116 cells, cells were seeded at 50% confluency 24 hours prior to labeling. For labeling, cells were incubated with 100 μM s^4^U for 24 hours, with medium replacement every 4 hours to maintain s^4^U activity. After pulse labeling, cells were washed three times with pre-warmed 1× PBS, and cultured in s^4^U-free medium containing 10 mM uridine to stop further labeling. Cells were harvested at 0, 2, 4, and 6 hours post-wash. Total RNA was extracted using TRIzol reagent and dissolved in nuclease-free water containing 1 mM DTT. *N. crassa* RNA samples were subjected to mRNA isolation using oligo(dT)25 magnetic beads (NEB, S1419S). For Thiol alkylation, a 50 μL reaction mixture containing 0.5-20 μg RNA, 5 μL iodoacetamide (IAA, 100 mM), 5 μL NaPO□ buffer (500 mM, pH 8.0), and 25 μL DMSO was incubated at 50°C for 15 minutes. The reaction was quenched by adding 1 μL of 1 M DTT. RNA was precipitated with 1 μL glycogen, 5 μL 3 M NaOAc, and 125 μL absolute ethanol at -80°C for 30 minutes. The pellet was resuspended in 5–10 μL nuclease-free water. For library preparation, employing the VAHTS Universal V8 RNA-seq Library Prep Kit (Vazyme, NRM605).

### RNA library and sequencing

To obtain RNA degradation profile of *N. crassa*, strain was subjected to same procedure as above described. Cultures were harvested at time points of 0, 20, 40, and 60 min after thiolutin treatment. For the subsequent steps, 15 μg of *N. crassa* total RNA and 150 ng of HEK293T RNA (served as a spike-in) were mixed together. The mRNA was isolated using oligo(dT)25 magnetic beads (NEB, S1419S) through two rounds of purification. Subsequently, 80 ng of purified mRNA was used for library preparation, employing the VAHTS Universal V8 RNA-seq Library Prep Kit (Vazyme, NRM605). For human RNA decay library, 1 μg total human RNA and 100 ng *N. crassa* total RNA (used as a spike-in) were input for library preparation. rRNA depletion was performed using de-ribo, with rRNA probes provided from Lai Lab ^78^. The library prepare was also performed with VAHTS Universal V8 RNA-seq Library Prep Kit. The inserted size of library was about 150-200 bp. The libraries were sequenced on MGI platform (MGISEQ-2000, BGI) and SURFSeq 5000 platform (GeneMind Biosciences Company Limited.).

### RNA decay profiling

Raw sequencing reads were adapter-trimmed and quality-filtered using Trim Galore (v1.3.0). Processed reads were aligned to combined *Homo sapiens* (GRCh38) and *Neurospora crassa* (NC12) reference genomes with STAR (v2.7.10b). Scale factors were calculated via multiBamSummary bins (deepTools v3.5.1) using spike-in alignments, then applied to normalize *N. crassa* or human cell sample read counts.

RNA half-lives were calculated per established methods ^79^ by: (1) normalizing raw counts to spike-in controls; (2) selecting genes exhibiting monotonic decay; (3) fitting exponential curves (*N(t)=N_0_* ⋅ *e ^−kt^*) to time-course data using Python (NumPy/SciPy’s curve_fit); and (4) deriving half-lives (*t_1/2_ =ln*(2)*/k*) representing 50% signal reduction from t=0. Biologically relevant half-lives (1-48 hours) were retained for analysis in human cell samples, with complete decay profiles cataloged in Table S10.

### SLAM-seq analysis

SLAM-seq data were processed through a custom computational workflow. Raw reads underwent adapter trimming and quality control using Trim Galore (v0.6.10), followed by alignment to reference genomes (Homo sapiensGRCh38 or *Neurospora* crassa NC12) via STAR (v2.7.10a) with default parameters to detect T>C conversions. Aligned reads were converted to sorted BAM files using SAMtools (v1.15.1). A custom Python script then isolated reads containing T>C mutations - indicative of 4-thiouridine incorporation - generating mutation-specific BAM files. Gene-level quantification was performed with featureCounts (Subread v2.0.3), separately enumerating: (1) T>C mutation-containing reads and (2) total mapped reads per gene (using genome-specific GTF annotations). The T>C mutation ratio (remaining reads percentage) was calculated as (T>C reads)/(total reads) for each gene. These ratios across timepoints were fitted to exponential decay curves (*N(t)=N_0_* ⋅ *e ^−kt^*) using SciPy’s curve_fit, with RNA half-lives derived from decay constants (*t_1/2_ =ln*(2)*/k*), representing time to 50% signal loss relative to t=0.

### Distribution of out-of-frame RF around HSC

Take the Ribo-seq A-site count bedGraph file as input. Define analysis windows spanning 31 or 61 codons. Slide these windows codon-by-codon across the transcriptome. Select windows centered on fixed-position HSC in either the +1 or -1 reading frame for standardization. For each qualifying window, calculating the average cumulative ribosome density at each codon position, and excluding fragments with <15 total A-site signals within the window. Then calculating frameshift rates by comparing ribosome densities in: +1 or -1 frame versus 0 frame at each codon position. Generate line plots showing frameshift rates across codon positions.

### Codon substitution and codon usage score plot

For removing hidden stop codons in test sequence, synonymous codon substitutions were performed to eliminate hidden stop codons while preserving the amino acid sequence. To maintain codon usage patterns comparable to wild-type (WT), substitutions employed synonymous codons with similar Relative Synonymous Codon Usage (RSCU) values. For codon optimization of selected fragments, all codons were substituted by the synonymous codon with the highest RSCU value, and new sequence without any HSCs. All modified sequences in this study were shown in Table S9.

Codon usage score plot was obtained by calculating the mean of RSCU value in 20 codons window size, and the window moved one codon distance to 3’ end after every calculation. The table of RSCU value was obtained from http://codonstatsdb.unr.edu/index.html/.

### Codon Bias Index (CBI) calculation

Using the codon statistics database website (http://codonstatsdb.unr.edu/index.html/), we determined the optimality of codons for each gene’s longest mRNA transcript in the organisms mentioned in this research.(Table S1) The CBI values were then calculated for each gene by using the formula described below:

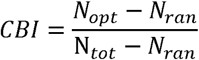

N_opt_ is the number of optimal codons in a gene, N_ran_ is the number of other codons within the gene except the optimal codon, N_tot_ is the number of all the codons within the gene. CBI value with more or less than 0 indicates a given gene with optimal or non-optimal codon usage preferring, the highest value (1) indicates all codons are optimal and 0 value is random codon usage.

### Codon Stabilization coefficient and codon off-frame translation coefficient

The codon stabilization coefficient (CSC) was calculated as previously established. ^8^ In brief, the longest CDS was chosen to obtain the codon frequency, and every gene with a half-life value was obtained from RNA degradation. The CSC was determined by calculating a Pearson correlation coefficient between the occurrence frequency of individual codons and the half-life of the mRNA harboring the codon.

For the calculation of codon out-of-frame translation coefficient (COC) in +1 and -1 frame, the out-of-frame ratio of each gene in ribosomal profiling was determined. Then the COC was determined by calculating a Pearson correlation coefficient between the frequency of occurrence of individual codons and out-frame ratio of the mRNAs harboring the codon.

### uORF identification

To identify genes containing translated uORF, RiboToolkit (https://rnainformatics.org.cn/RiboToolkit/) was employed for analysis. First, raw sequencing reads underwent adapter trimming and quality control using Trim-Galore (v0.6.10). The processed RPFs (Ribosome Protected Fragments) were then converted to FASTA format and uploaded to the RiboToolkit web server. Analysis was performed using default parameters, and genes with uORF translation events were subsequently identified based on the “Actively translated ORFs” output file. Additionally, pre-annotated uORF-containing genes for *Mus musculus* and *Homo sapiens* were downloaded from the database for validation.

### Correlation analysis

To evaluate the association between gene CBI and HSC density, the HSC density was determined by calculating the ratio of the number of termination codons in the +1 and -1 frames to the total length of each gene. Scatter plots were employed to illustrate the relationship between CBI and HSC density across all genes, and the correlation coefficient was computed to quantify the strength of this correlation.

In examining the correlation between RSCU and COC, scatter plots were produced using RSCU and COC values for 61 codons. And same analysis was performed to evaluating the correlation between CSC and RSCU, or between COC and CSC of 61 codons.

## QUANTIFICATION AND STATISTICAL ANALYSIS

Statistical analyses were conducted using GraphPad Prism and Python software. Three biological replicates were included in the analysis. For comparisons between two data sets, a Student’s *t*-test was performed, while for comparisons involving multiple data sets, a one-way ANOVA was applied. Statistical significance was considered valid with a *P* value of less than 10□□ for genome-wide analyses and less than 0.05 for regular analyses.

